# Non-neutralizing antibodies targeting the immunogenic regions of HIV-1 envelope reduce mucosal infection and virus burden in humanized mice

**DOI:** 10.1101/2021.05.24.445493

**Authors:** Catarina E. Hioe, Guangming Li, Xiaomei Liu, Ourania Tsahouridis, Xiuting He, Masaya Funaki, Jéromine Klingler, Alex F. Tang, Roya Feyznezhad, Daniel W. Heindel, Xiao-Hong Wang, David A. Spencer, Guangna Hu, Namita Satija, Jérémie Prévost, Andrés Finzi, Ann Hessell, Shixia Wang, Shan Lu, Benjamin K. Chen, Susan Zolla-Pazner, Chitra Upadhyay, Raymond Alvarez, Lishan Su

## Abstract

Antibodies are principal immune components elicited by vaccines to induce protection from microbial pathogens. In the Thai RV144 HIV-1 vaccine trial, vaccine efficacy was 31% and the sole primary correlate of reduced risk was shown to be vigorous antibody response targeting the V1V2 region of HIV-1 envelope. Antibodies against V3 also were inversely correlated with infection risk in subsets of vaccinees. Antibodies recognizing these regions, however, do not exhibit potent neutralizing activity. Therefore, we examined the antiviral potential of poorly neutralizing monoclonal antibodies (mAbs) against immunodominant V1V2 and V3 sites by passive administration of human mAbs to humanized mice engrafted with CD34+ hematopoietic stem cells, followed by mucosal challenge with an HIV-1 infectious molecular clone (IMC) expressing the envelope of a tier 2 resistant HIV-1 strain. Treatment with anti-V1V2 mAb 2158 or anti-V3 mAb 2219 did not prevent infection, but both reduced the virus burden, and V3 mAb 2219 displayed a superior potency compared to V1V2 mAb 2158. While these mAbs had no or weak neutralizing activity and elicited undetectable levels of antibody-dependent cellular cytotoxicity (ADCC), V3 mAb 2219 displayed a greater capacity to bind virus- and cell-associated HIV-1 envelope and to mediate antibody-dependent cellular phagocytosis (ADCP) and C1q complement binding as compared to V1V2 mAb 2158. Mutations in the Fc region of 2219 abolished these effector activities and abrogated virus control in humanized mice. These results demonstrate the importance of Fc functions other than ADCC for antibodies without potent neutralizing activity.

## Introduction

Almost forty years after the identification of HIV-1 as the virus that causes AIDS, ∼38 million people worldwide are living with the virus ^1^. Despite the achievement of effective virus suppression with combination antiretroviral therapies (cART) and improvements in prevention strategies that incorporate cART for treatment as prevention, pre-exposure prophylaxis, and post-exposure prophylaxis, 1.7 million new infections still occur in 2019, disproportionately affecting populations with limited access to care. Preventive vaccines would be powerful tools for ending this epidemic, but none are yet available and the development of HIV-1 vaccines has faced tremendous scientific challenges. To generate efficacious vaccines, a better understanding is required of protective immune components and functions.

Of the phase IIb/III HIV-1 vaccine efficacy trials, the Thai RV144 trial is the only one yielding a promising efficacy signal ^2^. Although the vaccine efficacy of 31% was modest, this trial provided the first indication of vaccine-induced immune correlates for protection against HIV-1 in humans. Among the six primary variables measured, high IgG levels against the V1V2 region of HIV-1 Env was identified to be a correlate of reduced HIV-1 acquisition risk ^3-5^. Subsequent studies defined additional correlates that include antibodies against the V3 loop in a subset of vaccine recipients with lower levels of Env-specific plasma IgA and neutralizing antibodies ^6,7^. Nonetheless, the mechanistic correlates for protection remain unclear. The RV144 vaccine-induced antibody responses did not display broad or potent virus-neutralizing activity and vaccine efficacy did not correlate with antibody-mediated neutralization. Instead, high levels of antibody-dependent cellular cytotoxicity (ADCC) in combination with lower plasma anti-Env IgA were detected among the secondary correlates ^3^. Many Env-specific monoclonal antibodies (mAbs) isolated from the RV144 trial participants also display ADCC activity ^8^, signifying the importance of non-neutralizing Fc-mediated functions of antibodies.

A previous study evaluating passive transfer of a non-neutralizing anti-gp41 mAb F240 in rhesus macaques demonstrated sterilizing protection in 2 of 5 animals and lower viremia in the remaining 3 animals after a high challenge dose of tier 2 SHIV SF162P3 ^9^. Protective activities were also observed in mice treated with non-neutralizing mAbs against West Nile ^10,11^ and influenza virus ^12-15^. In contrast, another passive transfer study with non-neutralizing anti-gp41 mAb 7B2 and SHIV BaL challenge showed no protection, although the number of transmitted/founder (T/F) variants was reduced ^16^. The administration of a non-neutralizing anti-gp41 mAb 246D to humanized mice with established HIV-1 infection also selected for escape mutation ^17^. In contrast, sterile protection was achieved by passive transfer of a V3-specific mAb KD247 into cynomolgous macaques challenged with SHIV strain C2/1, which was neutralized by KD247 at an IC50 value of 0.5 μg/ml ^18^. Administration of the V1V2-specific mAb 830A to rhesus macaques also was found to protect 5 of 18 animals that were repeatedly challenged with SHIV BaL and reduce plasma and cell-associated virus loads in blood and tissues of the remaining animals ^19^. However, V1V2 mAb 830A neutralizes the tier 1 SHIV BAL challenge virus with an IC50 value of 1.4 μg/ml, so whether neutralizing and/or non-neutralizing activities mediated the reduced infection was not clear. Until now, no studies have evaluated the in vivo efficacy of V1V2- and V3-specific antibodies with no or poor neutralizing activity against tier 2 viruses which represent the majority of HIV-1 isolates. Therefore, in this study we sought to evaluate their protective potential against HIV-1 by passive administration of non-neutralizing V1V2- and V3-specific mAbs to human CD34+ hematopoietic stem cell-engrafted mice capable of supporting HIV-1 infection.

In the present study, two human IgG1 mAbs, V1V2-specific 2158 and V3-specific 2219, were tested against an HIV-1 infectious molecular clone (IMC) with the tier 2 JRFL Env. MAb 2158 is specific for a conformation-dependent V2i epitope in the underbelly of the V1V2 domain near the integrin α4β7-binding motif ^20-23^. MAb 2219 recognizes the crown of the V3 loop by a cradle-binding mode ^24-26^. Both mAbs show a high degree of cross-reactivity with gp120 proteins from the major HIV-1 clades (A, B, C, D, F) and CRF02_AG, but the epitopes are masked in the functional Env spikes on most tier 2 HIV-1 virions, resulting in their inability to neutralize virus in the conventional in vitro assay ^22,27-31^. Indeed, neutralization screening against large arrays of HIV-1 pseudoviruses with tier 1-3 Envs from different clades showed that 2158 neutralizes only a few tier 1 strains and 2219 neutralizes <50% of tier 1 and tier 2 strains ^22,27,28^, even though pseudoviruses are more sensitive to neutralization than replication-competent viruses such as the JRFL IMC examined in this study. It is important to note, however, that unlike epitopes recognized by broadly neutralizing antibodies (bNAbs), the V2i and cradle V3 epitopes represent immunogenic sites that can be readily targeted by vaccination. This was evident from the detection of antibody responses against these specific epitopes in the vaccine recipients who participated in the VAX003, VAX004, and RV144 clinical trials, although durable responses remained unattainable ^5,7,32-35^.

In this study we examined plasma viremia and tissue-associated vRNA and vDNA burden in humanized mice that received V2i mAb 2158 or V3 mAb 2219 and challenged with JRFL IMC via the intrarectal route. We determined the importance of Fc-mediated functions by administering V3 mAb 2219 with Fc mutations that significantly decrease Fc receptor and/or complement binding to humanized mice to protect against JRFL IMC. The data demonstrate the contribution of antibody-dependent cellular phagocytosis (ADCP) and complement-dependent functions to suppress infection of neutralization-resistant HIV-1, thus providing an impetus for the development of vaccine strategies that harness these Fc-dependent antibody functions to control HIV-1 infection.

## Results

### Passively administered V2i mAb 2158 and V3 mAb 2219 display differential activities against rectal HIV-1 challenge in CD34+ HSC-engrafted humanized mice

To assess the protective potential of anti-HIV-1 mAbs with poor or no neutralizing activities, we passively administered V2i mAb 2158 and V3 mAb 2219 to CD34+ HSC-engrafted humanized NSG mice that were then challenged intrarectally with an HIV-1 vpr-deleted infectious molecular clone (IMC) expressing the tier 2 JRFL envelope ^36^. A third group of mice was given an irrelevant control mAb, 860, which is specific for the major capsid protein VP2 of parvovirus B19^37^. Mice (n=5/group, male and female) were given two doses of each mAb (700 µg/dose) intraperitoneally on day 0 and day 2, and challenged rectally with JRFL IMC at 2x 700 TCID_50_ at 4 and 24 hours following the first mAb dose (**Fig 1A**). The rectal challenge was chosen to represent a mucosal route pertinent for transmission to males and females. A total dose of 1400 TCID_50_/animal was determined by titration in a prior experiment to yield 100% infection in all exposed animals within a week. The mAb half-life was also assessed in the plasma following transfer to uninfected mice and found to be 11 days for V3 mAb 2219 and >14 days for V2i mAb 2158 following a plateau at 35 and 30 ug/ml for the respective mAbs (**Fig S1**).

**Figure 1:**
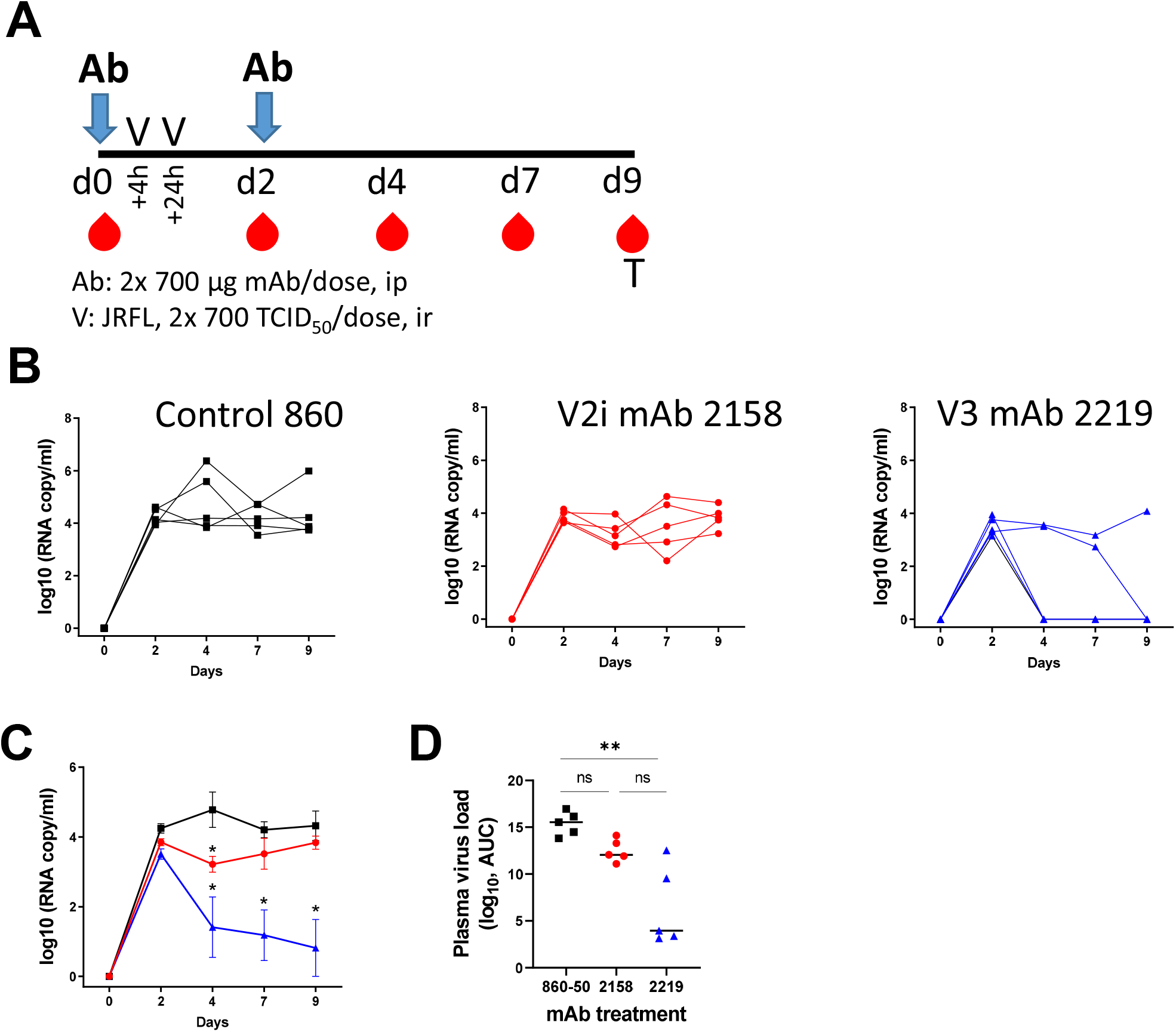
Passive transfer of V2i mAb 2158 and V3 mAb 2219 to CD34+ HSC‐engrafted humanized mice challenged with HIV‐1 JRFL IMC. A) Schematic of experimental protocol. B) Plasma vRNA loads from day 0 to day 9 in individual mice that received control mAb 860, V2i mAb 2158, or V3 mAb 2219 C) Mean plasma vRNA loads from day 0 to day 9 in each of the three groups. D) Area under the curve (AUC) of vRNA loads from days 0 to 9 in individual mice from each of the three groups. **, p<0.01 by Kruskal‐Wallis test with Dunn’s multiple comparison. ns, not significant (p >0.05).

Measurement of plasma vRNA loads showed that infection was established in all mice within 2 days of virus challenge (**Fig 1B**). In comparing the vRNA loads of individual mice receiving the control mAb 860 or V2i mAb 2158, the animals in both groups had similarly high viral loads from day 2 to day 9 (**Fig 1B**). Nonetheless, a brief but significant drop in the average vRNA load was observed on day 4 in the V2i mAb 2158-treated animals compared to the mAb 860 control group (**Fig 1C**). In addition, the overall virus burden of the 2158-treated group, as measured by area under the curve (AUC) of vRNA copies from days 0 to 9, tended to be lower than that of the control 860 group (**Fig 1D**). In contrast, the recipients of V3 mAb 2219 displayed a significant reduction in vRNA loads at days 4, 7, and 9 as compared to the 860 control group, with three of the five 2219-treated animals having viral loads below detection (**Fig 1B-C**). Moreover, a significant reduction in the vRNA AUC from day 0 to day 9 was observed in 2219-vs 860-treated groups (**Fig 1D**).

After termination of the experiment on day 9, the spleen, bone marrow, and lymph nodes were collected to quantify the levels of cell-associated vRNA, vDNA, and p24. As expected, vRNA and vDNA levels in the spleen, bone marrow, and mesenteric lymph nodes of the V3 mAb-treated 2219 group were low to undetectable, in comparison to the control 860 group (**Fig 2A-B**). The three animals with undetectable plasma viral loads on days 4-9 post infection also had undetectable tissue-associated vRNA and vDNA levels. In the V2i mAb 2158 group, we also observed significantly lower levels of cell-associated vRNA and vDNA in most tissues as compared to the control 860 group (**Fig 2A-B**). However, the reduction in the levels of vDNA in the lymph nodes of both 2158- and 2219-treated groups did not reach statistical significance (**Fig 2B**). The data demonstrate that although the passive transfer of V3 mAb 2219 and V2i mAb 2158 failed to prevent the establishment of virus infection, both mAbs showed the capacity to decrease virus burden. The V3 mAb 2219 exhibited a superior potency to control HIV-1 infection in vivo compared to V2i mAb 2158, suggesting a differential capacity to mediate effector functions.

**Figure 2:**
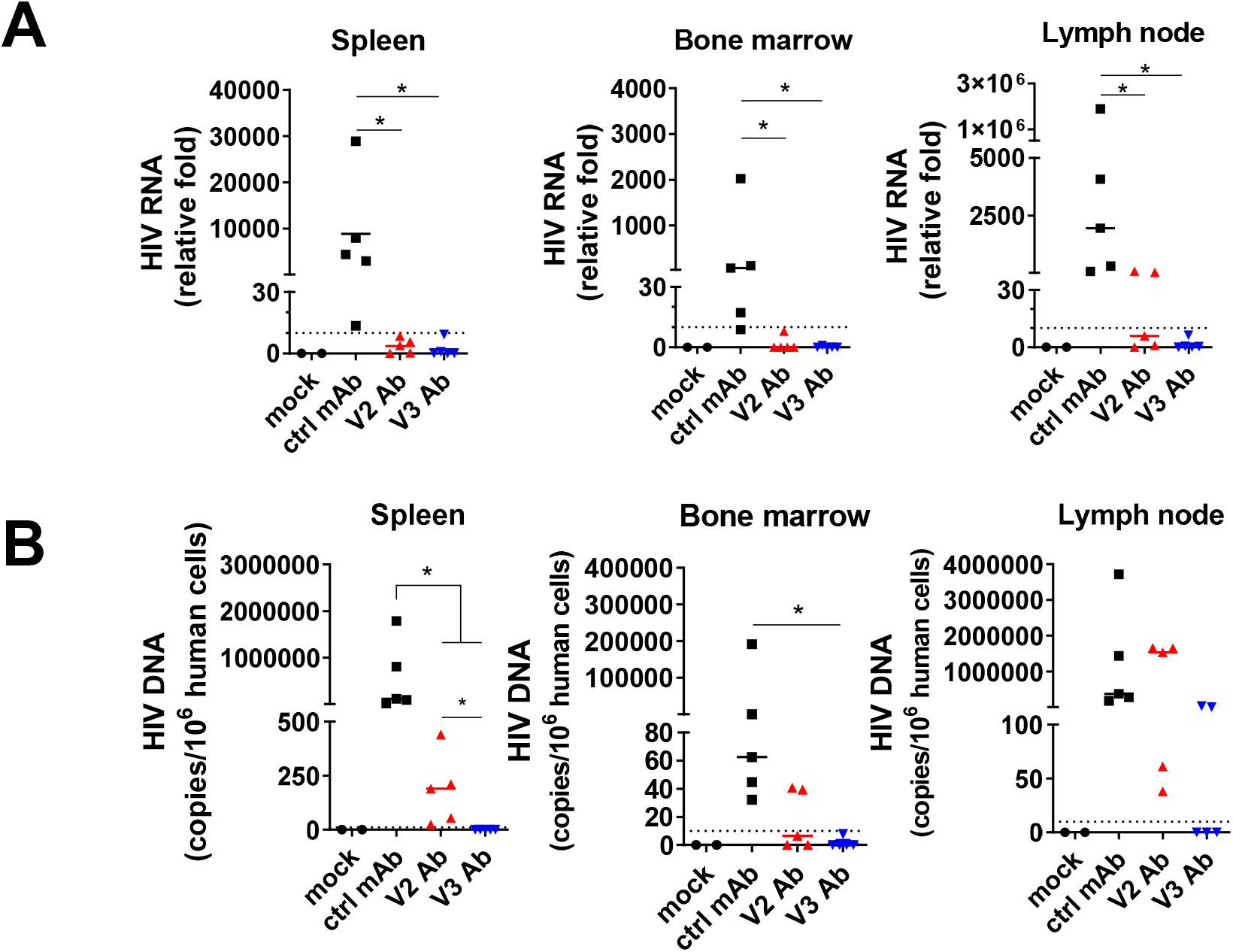
Levels of cell‐associated vRNA, vDNA, and Gag p24 in CD34+ engrafted humanized mice treated with mAbs and infected with JRFL IMC. A) Relative levels of cell‐associated vRNA in spleen, bone marrow, and mesenteric lymph nodes. B) Relative levels of vDNA in spleen, bone marrow, and mesenteric lymph nodes. ^*^, p <0.05 by Kruskal‐Wallis test with Dunn’s multiple comparison.

### V3 mAb 2219 binds better to virion- and cell-associated Env than V2i mAb 2158

Next, the V3 mAb 2219 and V2i mAb 2158 were examined for the ability to recognize different forms of HIV-1 Env. Both mAbs were isolated from US subjects infected with clade B viruses ^27^. 2219 and 2158 displayed strong ELISA reactivity with recombinant soluble gp120 proteins of JRFL and several other strains from different HIV-1 clades (**Fig 3A-C**). The EC50 titers of 2219 were 1.0-to 2.9-fold greater than 2158, depending on the HIV-1 gp120 strains tested (**Fig 3C**). The kinetics analysis by biolayer interferometry further showed that while V3 mAb 2219 and V21 mAb 2158 have similar KD values in the picomolar range for recombinant JRFL gp120, V3 mAb 2219 has a 14-fold faster Kon rate compared to V2i mAb 2158 (**Fig 3B**). Interestingly, the V3 mAb 2219 also demonstrated as much as 8-fold greater binding to virion-derived solubilized Env from JRFL IMC than V2i mAb 2158 (**Fig 3D-E**), although the relative binding to other solubilized Envs ranged from 5-fold weaker for REJO to 18-fold stronger for BG505 (**Fig 3E**). In these ELISA assays, recombinant gp120 proteins were directly coated onto ELISA plates (**Fig 3A-B**), whereas solubilized Env in virus lysate was captured by ConA coated on the plates (**Fig 3D-E**) before incubation with 2219 or 2158.

**Figure 3:**
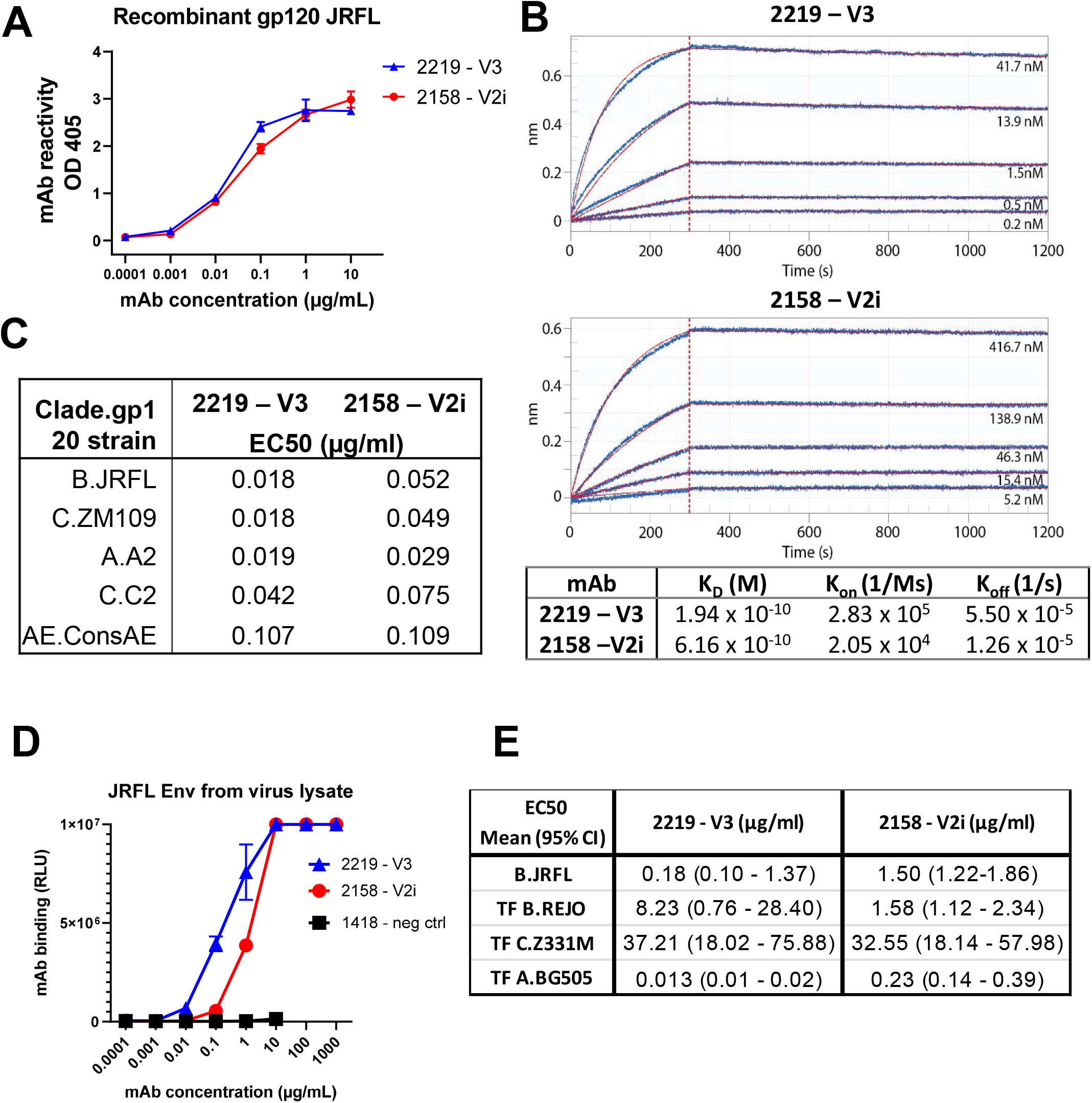
The binding strength of V2i vs V3 mAb to recombinant JRFL gp120 or virus‐derived gp120. A) Direct ELISA measurement of mAb reactivity to recombinant JRFL gp120 (1 µg/ml coated on plates) B) Kinetics analysis of mAbs binding to recombinant JRFL gp120 by biolayer interferometry. Fitted curves (1:1 binding model) are shown in red. Molar concentrations of gp120 are indicated. C) EC50 values of mAb binding to recombinant gp120 proteins from different HIV‐1 strains D) Measurement of mAb binding to gp120 from virus‐derived solubilized JRFL Env captured with ConA in sandwich ELISA. E) EC50 values of mAb binding to Env derived from JRFL and other HIV‐1 strains. HIV‐1 clade or CRF is indicated by a letter before the strain name. TF: transmitted founder.

Having compared the HIV-1 Env binding of 2219 and 2158 in ELISA-based assays, we investigated their abilities to bind Env on virion and cell surfaces. The differences between 2219 and 2158 were more evident in their ability to bind virion- and cell-associated Env (**Fig 4**). As compared to 2158, 2219 was observed to capture significantly higher levels of free JRFL virions (**Fig 4A-B**). Virus capture was measured after virus-mAb incubation for 24 hours. 2219 also bound cell surface-expressed JRFL Env and CD4+ CCR5+ CEM.NKr cells treated with recombinant JRFL gp120, similar to other V3 mAbs 2557 and 391/95 (**Fig 4C-D**). In contrast, 2158 recognized neither JRFL Env expressed on cell surface of transfected 293T cells nor JRFL gp120 tethered on the surface of CD4+ cells (**Fig 4C-D**). Higher levels of 2219 vs 2158 mAb binding were also detected using a Jurkat cell system previously used to characterize the binding profiles of anti-HIV-1 IgG monoclonal and polyclonal Abs that bind native-like HIV-1 Env^38^. Using this system, we compared the levels of mAb binding to JRFL, NL4.3, and three clade B transmitted founder virus Envs (QH0692; WITO4160; RHPA4259) (Fig 4E). For these experiments, 2219 and 2158 were compared with a parvovirus-specific control mAb (1418), another V3-binding mAb (447-52D), and the CD4 binding site targeting mAb b12. We observed that 2219 bound to all of HIV-1 envelopes tested except NL43, and displayed a similar binding pattern to the other V3 mAb tested, 447-52D (**Fig 4E**). In contrast, 2158 recognized JRFL at detectable but lower level than 2219. For most of the HIV-1 Envs tested, the level of 2158 binding was comparable to that of the irrelevant anti-parvovirus negative control mAb 1418 (**Fig 4E**).

**Figure 4:**
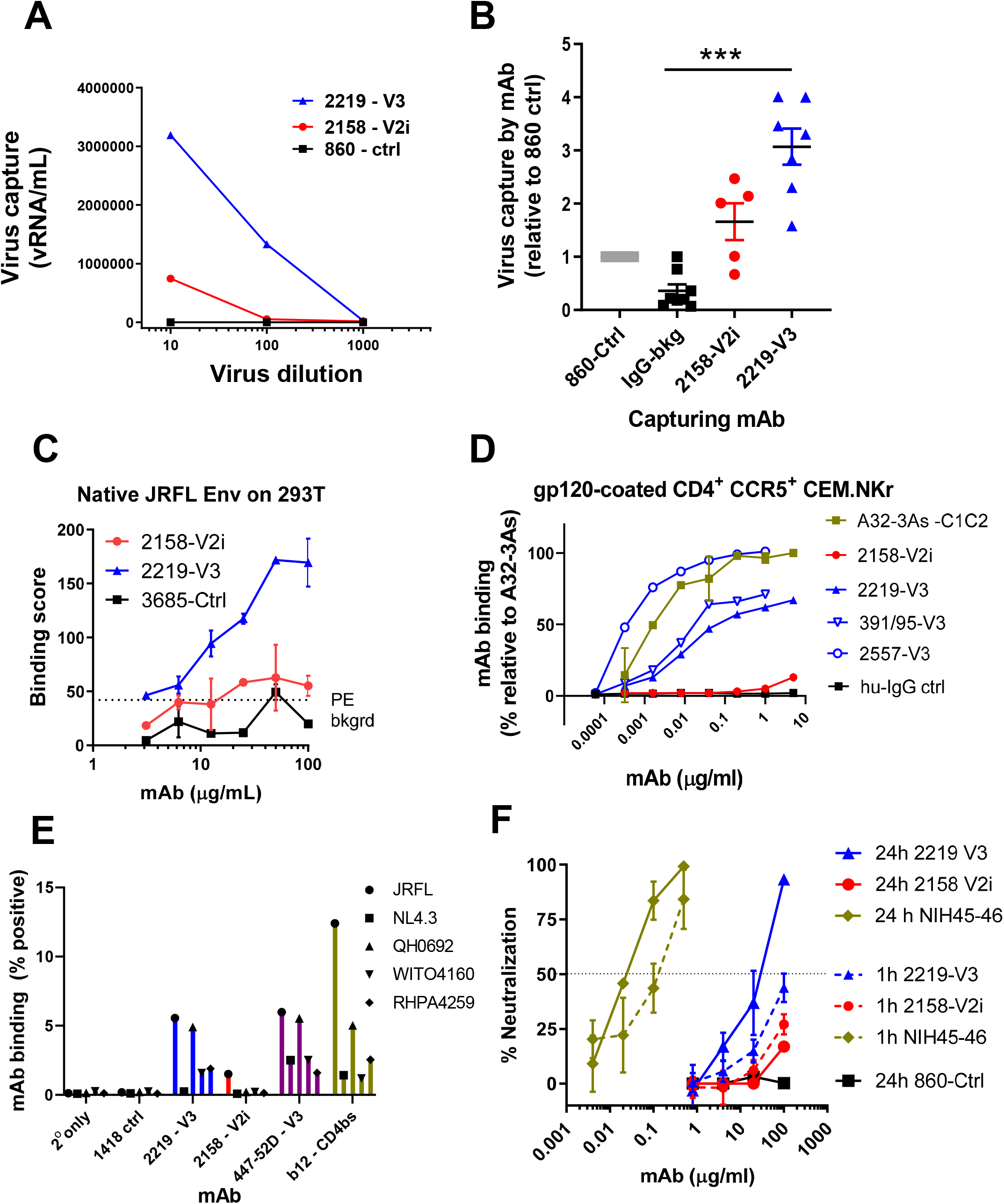
Capacity of V2i mAb 2158 and V3 mAb to bind and neutralize virus. A) Capture of JRFL IMC virions by 2158 vs 2219 with titrated amounts of virus input. Irrelevant mAb 860 was used as negative control. Data from one representative experiment. B) Capture of JRFL IMC virions by 2158 vs 2219 at a fixed virus input (5 to 6.5 log10 vRNA copies). Purified human HIV‐seronegative IgG and mAb 860 were used as negative controls. Virus capture was calculated relative that of control mAb 860 (set to 1). Data from 2‐3 experiments are shown. ***, p <0.001 by Kruskal‐Wallis test with Dunn’s multiple comparison. C) Binding of 2158 vs 2219 to native JRFL Env on transfected 293T cells. Irrelevant control mAb 3685 was used as negative control. D) Binding of 2158 vs 2219 to recombinant gp120‐coated CD4^+^ CEM.NKr cells. V3 mAbs 391/95 and 2557 were tested for comparison. MAb A32‐3As and human IgG were used as positive and negative controls, respectively. The relative level of mAb binding was calculated based on the binding of A32‐ 3As at 10 µg/ml (set at 100%). E) MAb binding to JRFL and other HIV‐1 strains (all clade B) produced and tethered on Jurkat cells. MAbs 2158 and 2219 were compared with V3 mAb 447 and CD4bs mAb b12. HIV‐1 Δvpu constructs bearing an mCherry reporter gene were used to transfect tetherin^hi+^ Jurkat cells. F) Neutralization of JRFL IMC by 2158 vs 2219 after virus‐mAb preincubation for 1 hour or 24 hours using TZM.bl target cells.

### V3 mAb 2219, but not V2i mAb 2158, exhibits weak and delayed virus-neutralizing activity

The V3 crown and V2i mAb epitopes are often occluded in functional Env trimers of most HIV-1 strains. However, due to Env structural flexibility, access to these epitopes can occur following extended incubation time to result in virus neutralization ^39,40^. When we tested 2219 and 2158 against JRFL IMC, no neutralization was observed for either mAb following a 1-hour virus-mAb incubation (**Fig 4F**). However, when the incubation time was extended to 24 hours, >50% neutralization was achieved by 2219. The neutralizing potency was >3 log10 weaker than the CD4-binding site (CD4bs)-specific bNAb NIH45-46 tested under the same condition. With 2158, virus neutralization was not achieved even after a 24-hour incubation. Hence, unlike V2i mAb 2158, V3 mAb 2219 had detectable, albeit delayed, neutralizing activity against JRFL IMC. This result is in accord with the greater Fab-mediated capacity of 2219 as demonstrated by virion capture and Env binding on the cell surface (**Fig 4A-E**). Notably, a concentration of 30 µg/ml, the equivalent to the IC50 value of 2219, was maintained in plasma over the nine days in which the animals were followed for viremia control (**Fig 1 and S1**). Nonetheless, in view of minimal neutralization capacity, the non-neutralizing activities are likely to play a role in the virus control observed with each of these mAbs in the humanized mouse experiments.

### V3 mAb 2219 mediates higher levels of Fc⍰IIa activation, ADCP, and C1q binding as compared to V2i mAb 2158

To determine the Fc-dependent functions mediated by V3 mAb 2219 and V2i mAb 2158, we first measured the capacity of these mAbs to activate Fc⍰RIIa and induce ADCP. Fc⍰RIIa is the primary Fc-receptor that activates ADCP in monocytes and macrophages in response to IgG-opsonized antigens ^41^. We employed an Fc⍰RIIa signaling assay with JRFL Δvpu that has previously been used to quantify differences in FcgR activation in HIV-1 controllers ^38^. Using this assay, we detected a higher capacity of 2219 to induce Fc⍰RIIa-mediated signaling as compared to 2158 (**Fig 5A**). To assess the ADCP activity of these mAbs, we used gp120-coated beads and Fc⍰RIIa+ THP-1 cells as phagocytes ^32,42^. In line with the Fc⍰RIIa signaling assay, we detected higher levels of ADCP activity mediated by 2219, as compared to 2158. However, both 2219 and 2158 had ADCP activity above the negative control mAb.

**Figure 5:**
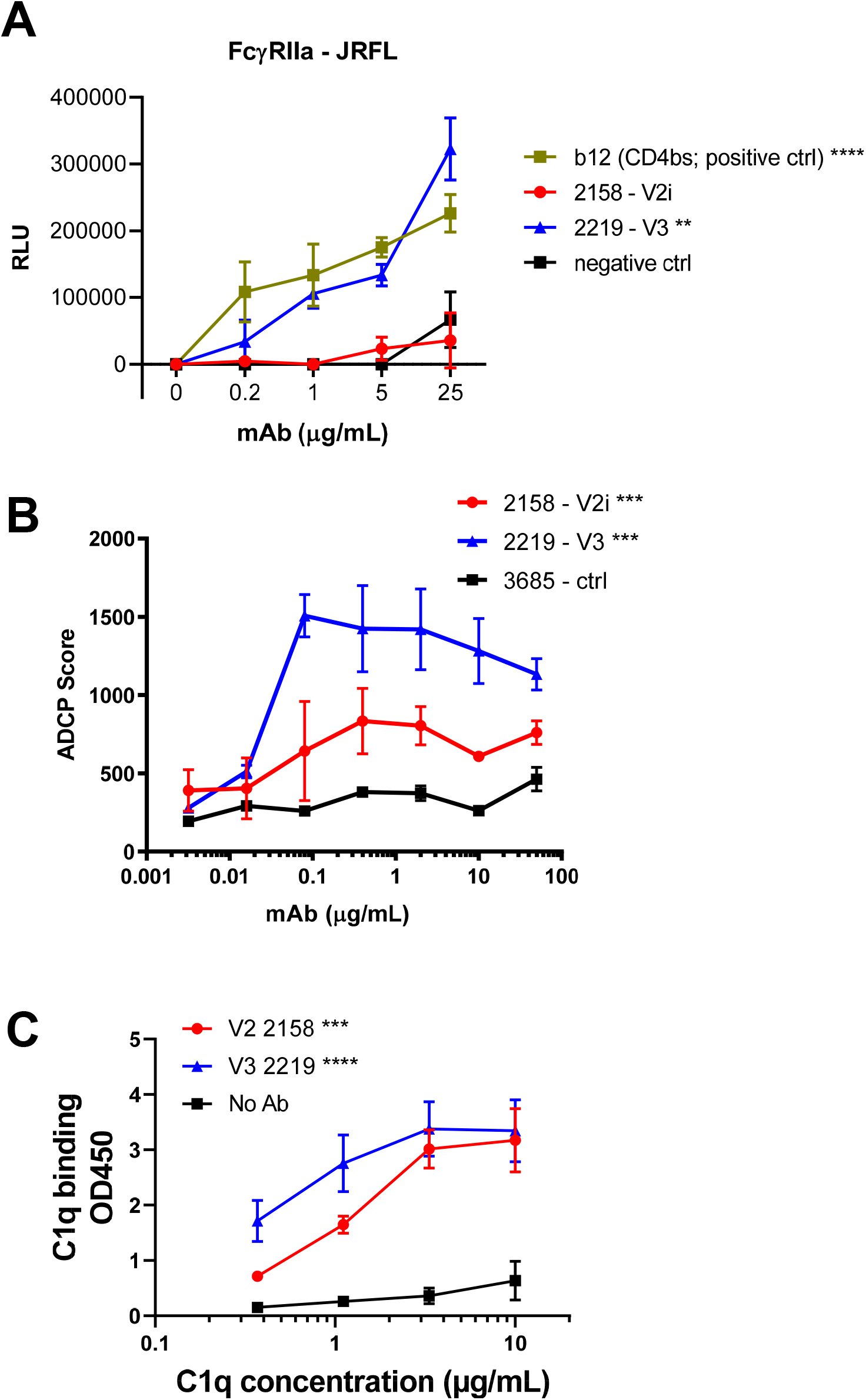
FcγRIIa signaling, ADCP, and complement binding activities of V2i mAb 2158 vs V3 mAb 2219. A) FcγRIIA signaling was measured by co‐incubating JRFL Δvpu‐nucleofected Jurkat cells with Jur‐γRIIa luciferase reporter cells in the presence of V2i, V3, or control mAbs. CD4‐binding site mAb b12 served as a positive control. RLU: relative light unit. ****, p <0.0001; **, p <0.01 by two‐way ANOVA vs 2158 and negative control. B) ADCP activity was measured using gp120 ZM109‐coated beads and THP‐1 phagocytic cells in the presence of titrated amounts of mAbs. Mean and SEM from 2 repeat experiments are shown. ***, p <0.001 by two‐way ANOVA for 2158 vs 2219 and for 2158 and 2219 vs negative control. B) C1q binding was measured using an ELISA‐based assay in which mAbs were reacted with gp120 on ELISA plates and then treated with titrated concentrations of C1q. C1q binding was detected using anti‐C1q antibodies and an alkaline phosphate‐conjugated secondary antibody. Mean and SD of replicate wells from a representative experiment are shown. **** p <0.0001; ^***^ p <0.001 by two‐way ANOVA vs no Ab control.

We also evaluated complement binding to immune complexes made with gp120 and 2219. For this, C1q deposition was measured as the first step in the classical mAb-dependent complement cascade. Both immune complexes made with 2219 and with 2158 showed dose dependent C1q deposition (**Fig 5C**). A slightly higher C1q binding activity was observed with 2219 vs 2158 in agreement with the gp120-binding EC50 titers of these mAbs (**Fig 3A-B**).

### V3 mAb 2219 and V2i mAb 2158 lack the ability to activate FclllRIIIa signaling and induce ADCC

We subsequently examined ADCC, an Fc-dependent activity that also has been shown to play a role in controlling viral infection where NK cells primarily mediate the killing of virally infected cells ^43^. ADCC is initiated when an effector cell expressing Fc⍰RIIIa engages with an infected target cell that has been opsonized with virus-specific IgG. To examine the ability of 2219 and 2158 to activate Fc⍰RIIIa, we used an Fc⍰RIIIa signaling assay with JRFL Δvpu and observed that both 2219 and 2158 failed to induce any significant Fc⍰RIIIa signaling in response to virus-infected target cells (**Fig S2 A**). In contrast, the CD4bs mAb, b12, efficiently induced Fc⍰RIIIa activation in an Ab-dependent manner in response to virus-infected target cells.

To investigate these findings further, we examined the capacity of 2219 and 2158 to induce ADCC using three different assay systems. These assays provide the opportunity to examine ADCC activity in the context of three distinct pairs of Env-bearing target cells and effector cells. As expected for V2i mAb 2158 that had no binding activity to cell-associated gp120 or full length Env **(Fig 4C-E**), no ADCC activity was detected in each of the three different assays (**Fig S2 B-D**). However, V3 mAb 2219 also displayed no detectable ADCC against JRFL gp120-coated CEM-NKr-CCR5 cells (**Fig S2 B**), even though binding to these target cells was readily detected (**Fig 4D**). The same result was observed with two other V3 mAbs: cradle-type 2557, similar to 2219 and ladle-type 391/95. MAbs 2219 and 2158 also failed to mediate ADCC against SHIV-SF162P3-infected NKR24 reporter cells (**Fig S2 C**). In the third assay that utilizes full-length JRFL IMC-infected primary CD4 T cells, 2219 again showed no binding and no ADCC (**Fig S2 D**).

Since the HIV-1 accessory proteins Vpu and Nef are known to impede ADCC responses by controlling Env accumulation at the surface of infected cells and limiting the Env-CD4 interaction that exposes the V3 and other CD4-induced (CD4i) epitopes ^44^, we examined JRFL IMC lacking Vpu and Nef for comparison. Similar to all other V3 mAbs tested (19b, GE2-JG8, 2424, 2557, 3074, 447-52D, and 268D), 2219 recognized JRFL IMC-infected cells and exhibited ADCC activity when Nef and Vpu were deleted (**Fig S2 D**). In contrast, the binding and ADCC activity of 2158 was not substantially improved by Nef and Vpu deletions, indicating differential mechanisms regulating the exposure of V2i vs V3 epitopes as previously reported ^40,45^.

### Significant contribution of V3 mAb 2219 Fc functions against rectal HIV-1 challenge in CD34+ HSC-engrafted humanized mice

Since V3 mAb 2219 showed the capacity to mediate ADCP and complement binding, we sought to examine the contribution of each of these Fc functions in controlling HIV-1 infection in vivo by introducing Fc mutations in the 2219 mAb. A double LALA mutation (L234A/L235A) was made to abolish ADCP and complement activation ^46^. Indeed, these Fc changes resulted in significant reduction of ADCP in an assay using the THP-1 phagocytic cells (**Fig 6A**) and complement activation as measured by C1q and C3d deposition (**Fig 6B**). The single KA mutation (K322A), on the other hand, was generated to abrogate C1q and C3d binding without affecting ADCP (**Fig 6A-B**) ^46^. As expected, we also observed a reduction in ADCP activity with 2219 LALA variant, as compared to the 2219 KA variant and 2219 WT, using an ADCP assay with mouse resident peritoneal mononuclear cells (**Fig 6C**). The LALA and KA mutations had no effect on the Fab-dependent capacity of 2219, as measured by ELISA reactivity with JRFL gp120 (**Fig S3 A**) and neutralization of JRFL IMC following a 24-hour virus-mAb incubation (**Fig S3 B**).

**Figure 6:**
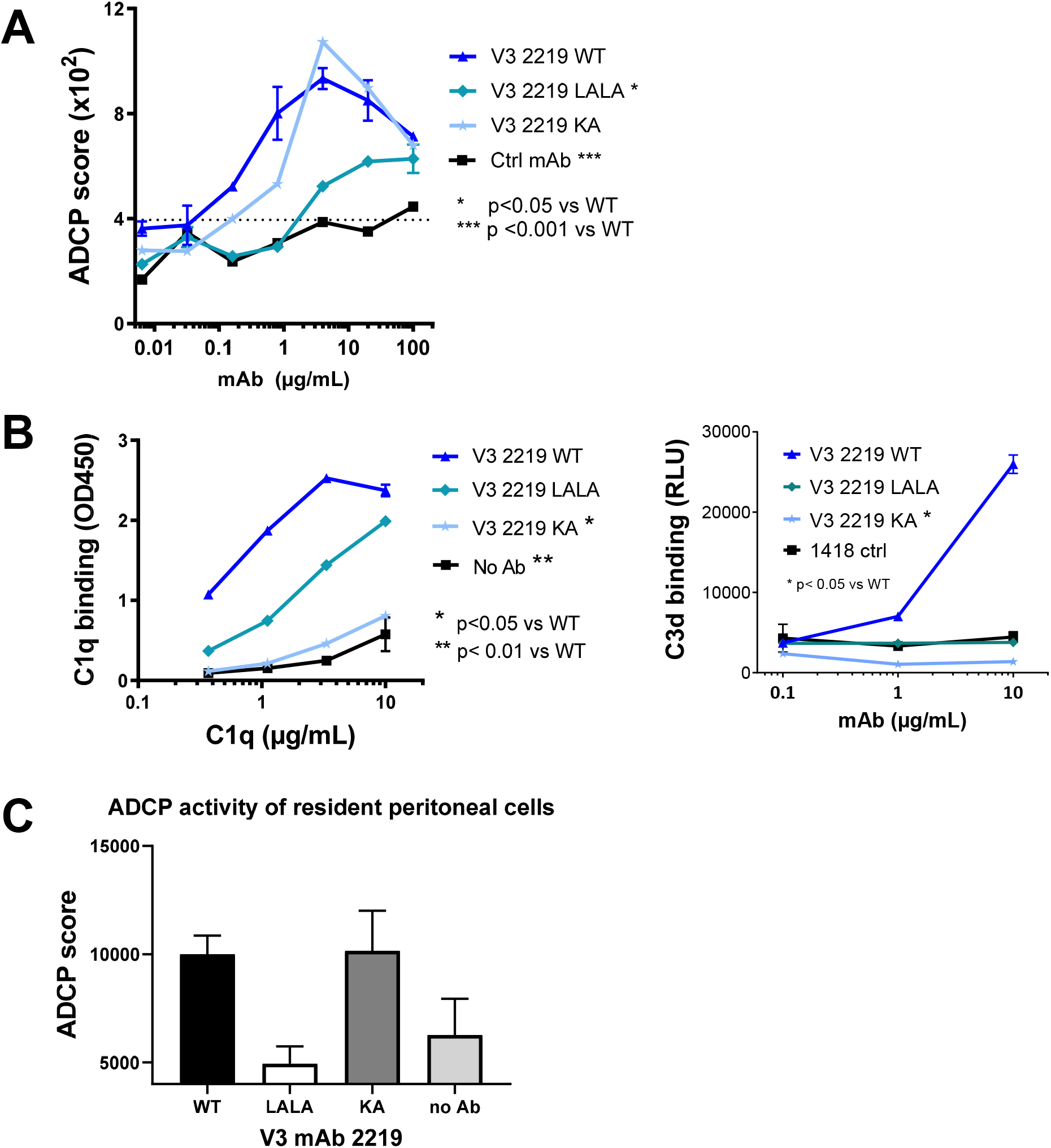
ADCP and complement binding activity of V3 mAb 2219 with WT or mutated Fc fragments (LALA and KA). A) ADCP activity was measured using gp120 ZM109‐coated beads and THP‐1 phagocytic cells in the presence of 2219 WT or Fc mutants. Statistical test was done using two‐way ANOVA. B) Complement binding by V3 mAb 2219 with WT and mutated Fc fragments was detected in ELISA or multiplex bead experiments in which C1q or C3d deposition to mAbs in complex with V3 MN peptide was measured with anti‐C1q or anti‐C3d secondary antibodies. Comparison was analyzed with two‐way ANOVA. C) ADCP of 2219 WT and Fc mutants was detected by measuring phagocytosis of mAb‐treated gp120 ZM109‐coated beads by resident peritoneal macrophages from NSG mice.

To examine the in vivo effects of these Fc mutations, the 2219 LALA and KA variants were administered intraperitoneally to CD34+ HSC-engrafted mice. The wild type 2219 and irrelevant control mAb 860 were tested in parallel for comparison. The 4 groups of mice (5/group) were given mAb intraperitoneally on day 0 and rectally challenged with JRFL IMC 4 hours later (**Fig 7A**). The mAb and virus treatment was repeated on days 2-3. Each animal received 2 x 700 ug mAb and was challenged with 2x 700 TCID50 of JRFL IMC. Plasma virus loads were measured starting from day 2 up to day 14. Data in **Fig 7B-C**.show that the 2219 WT group had lower plasma virus burden as compared to the control group as measured by virus load AUC values, and that the blunting effect was most apparent on the virus peak at day 2. Animals in the LALA and KA groups also showed dampened peak plasma vRNA loads, but the virus load AUC values were not significantly different from the control group. Similar results were observed when these mAbs were tested on another cohort of mice generated with CD34+ HSCs from a different donor (**Fig S4**).

**Figure 7:**
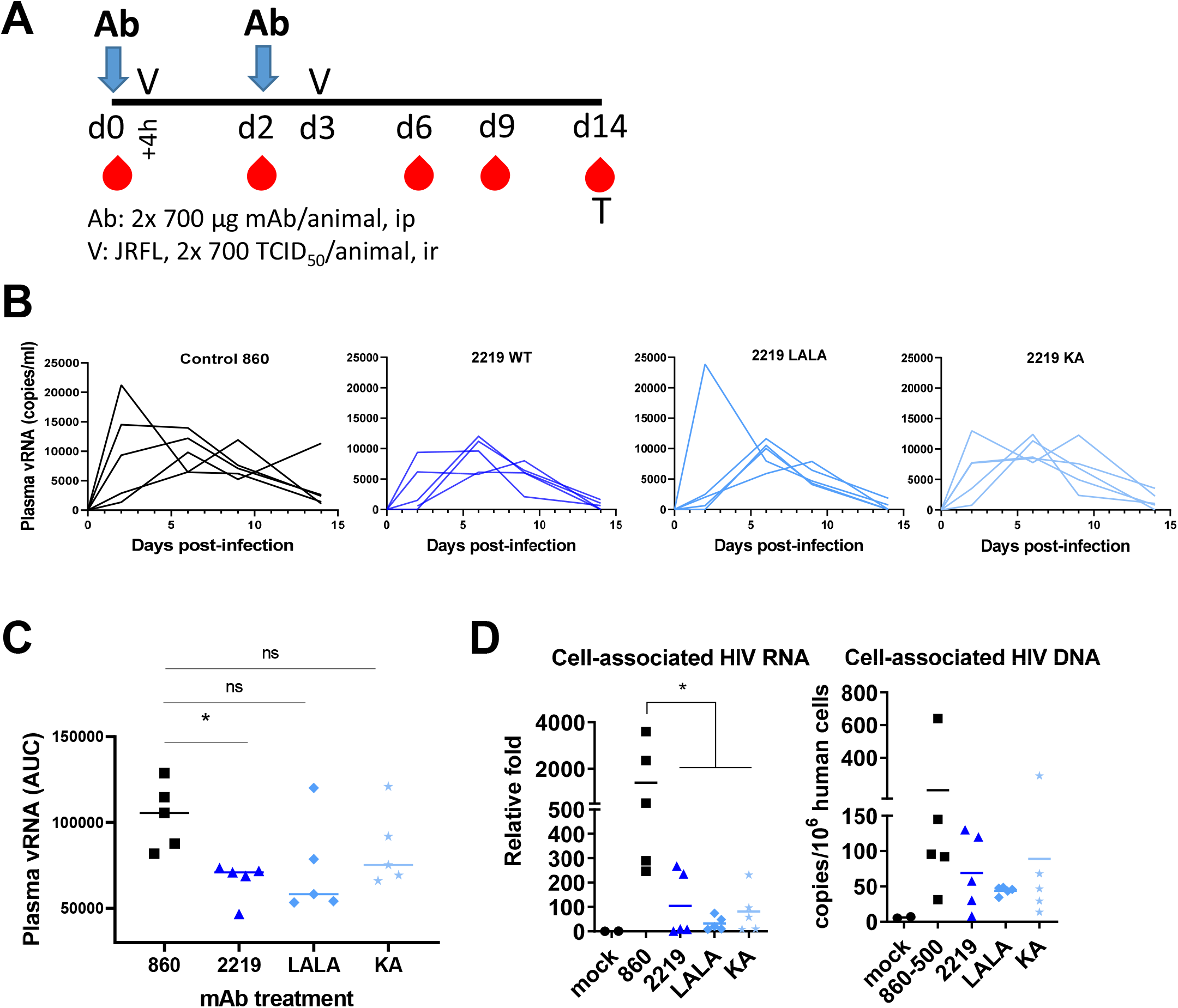
Virus control by passively transferred 2219 WT vs LALA vs KA in humanized mice challenged with JRFL IMC. A. Schematic of experimental protocol B. Plasma viremia levels in individual mice from day 0 to day 14. C. AUCs of vRNA in individual mice from day 0 to day 14. D. Relative levels of cell‐associated vRNA and vDNA in the spleen on day 14. ^*^, p <0.05 by Kruskal‐Wallis test with Dunn’s multiple comparison.

When the splenocytes were collected on day 14, we observed reduced levels of cell-associated vRNA in the WT, LALA, and KA groups vs the control 860 group (**Fig 7D**). However, vDNA remained unchanged (**Fig 7D**). Measurement of p24+ cells further revealed that lower percentages of p24+ cells were found among human CD45+ CD3+ CD8-T cells in the spleen of mice that received 2219 WT vs control 860 (**Fig 8A**). The percentages of p24+ cells in the LALA and KA groups, on the other hand, were higher than that of the 2219 WT group. These data demonstrate that LALA and KA mutations diminished virus suppressive activity of 2219. Together with data in **Fig 7**, the results indicate the important contribution of Fc-dependent functions, in particular complement activation, to the antiviral potency of V3 mAb 2219.

**Figure 8.**
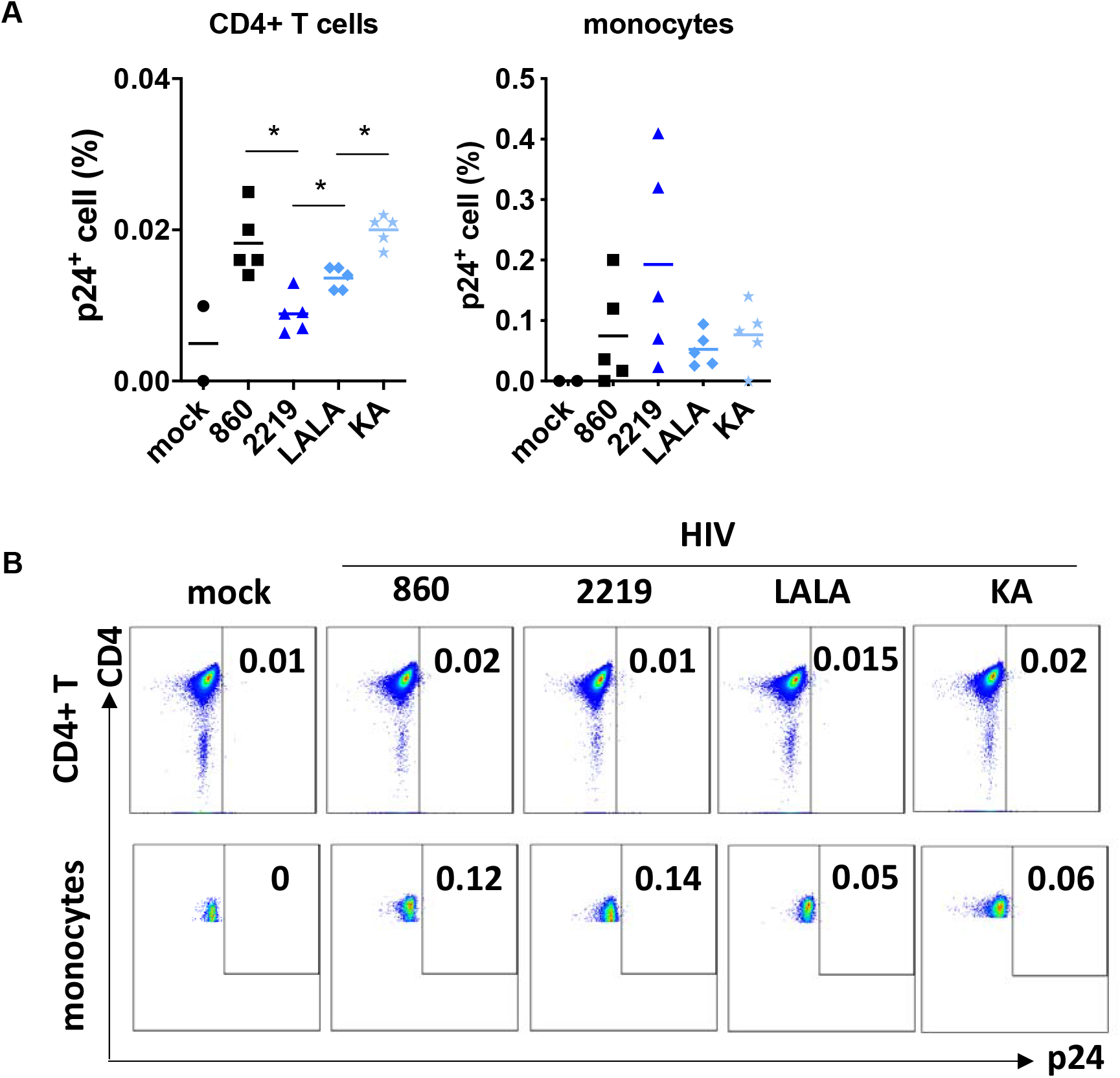
Detection of p24+ cells in the spleen of humanized mice treated with 2219 WT or Fc mutants and challenged with JRFL IMC. Spleen cells were collected from each animals shown in Figure 7 and subjected to intracellular staining with anti‐p24 mAb KC57 and staining for markers of cell viability, human CD45 (huCD45), mouse CD45 (mCD45), CD4 T cells (CD3+CD8‐), and monocytes (CD3‐CD11c‐CD14+). Analysis was done on viable human cells gated as hCD45+ and mCD45‐. A. Percentages of p24+ cells among CD4 T cells and monocytes in animals treated with 860 (irrelevant mAb), 2219 WT, 2219 LALA, or 2219 KA, and challenged with JRFL IMC. Cells from mock‐infected mice were included as negative controls. ^*^, p <0.05 by ANOVA with multiple comparison test. B. Dot plots from representative animals in the treated and mock groups.

We noted that although the percentages of p24+ CD4 T cells in the 2219 WT group were reduced to almost the mock background (**Fig 8A**), plasma virus loads were not fully suppressed (**Fig 7B-C**), suggesting the involvement of other cell types supporting virus infection. We examined p24+ cells among human CD45+ populations of CD4 T cells (CD3+ CD8) and monocytes (CD3-CD11c-CD14+). In the control mice, p24+ cells were detected at varying percentages in CD4 T cells and monocytes (**Fig 8A-B**). In the 2219 WT group, while lower percentages of p24+ cells were seen among CD4 T cells as described above, no decrease was measurable in monocytes. Similarly, no reduction of p24+ cells was seen in monocytes of the LALA and KA groups. These data point to the disparities in the ability of V3 mAb 2219 to control virus infection in different cell types and the lack of antiviral potency against virus reservoirs beyond CD4 T cells.

## Discussion

This paper provides the first evidence for the ability of passively infused V2i mAb 2158 and V3 mAb 2219 to reduce levels of cell-associated vRNA and vDNA in a CD34+ HSC-engrafted humanized mouse model upon challenge with a resistant tier 2 HIV-1, JRFL IMC. Neither mAbs showed detectable neutralizing activity against the challenge virus in the standard TZM.bl neutralization assay, but both mAbs could mediate Fc-dependent antiviral activities. Notably, V3 mAb 2219 displayed a greater capacity than V2i mAb 2158 to bind free virions, cell-associated virions, cell-bound gp120, and membrane-associated Env on transfected or infected cells. Unlike V2i mAb 2158, V2 mAb 2219 was able to exert delayed neutralization against JRFL IMC detectable after prolonged mAb-virus pre-incubation, although the IC50 titer was >3 log less potent than that of bNAb such as NIH45-46. Correspondingly, V3 mAb 2219 exhibited a greater control of virus infection in humanized mice, as indicated with reduced levels of vRNA in plasma and cell-associated vRNA and vDNA in tissues. V2i mAb 2158, on the other hand, lowered cell-associated virus burden in tissues with minimal effects on plasma viremia. In the absence of potent neutralizing activity, these results suggest a significant involvement of non-neutralizing functions in the anti-HIV-1 suppressive mechanisms wielded by these mAbs.

Both V2i mAb 2158 and V3 mAb 2219 showed the capacity to activate Fc⍰RIIa signaling and induce Fc⍰RIIa+ THP-1 phagocytic cells to mediate ADCP. Upon immune complex formation, both mAbs also were able to engage C1q, the initial step in the classical complement cascade. The Fc⍰RIIa- and complement-mediated functions were abrogated by the introduction of Fc mutations, and these mutations reversed virus suppression observed in the wild type 2219-treated humanized mice. Of note, the KA mutation that only abrogated complement binding had the same effect as the LALA mutations that decreased both ADCP and complement activation, indicating the importance of complement-mediated functions in HIV-1 control by V3 mAb 2219. Nonetheless, 2219 was not able to suppress infection completely.

In contrast to their capacity to trigger Fc⍰RIIa signaling and mediate ADCP, neither 2158 nor 2219 had measurable ADCC activity in all 3 assays performed with different target and effector cells. These results are consistent with past data showing that the ADCC activity detected in a modified rapid fluorescent ADCC assay with gp140-treated CEM-NKr cells and NK effector cells was <10% for both mAbs ^47^. In an ADCC assay that utilized CD4 T cells infected with JRFL IMC as target cells vs the Nef- and Vpu-deleted counterpart, the failure of V3 mAb 2219 to recognize and elicit ADCC against JRFL IMC-infected target cells was overturned by Nef and Vpu deletion. Among the manifold effects of Nef, the downregulation of CD4 has been attributed to minimizing CD4-induced epitope exposure, while the reduction of NKG2D ligand expression increases the resistance of HIV-1-infected cells to ADCC ^48-51^. Apart from contributing to CD4 downregulation, Vpu promotes virus release from infected cells by counteracting the restriction factor tetherin (BST2), thus limiting accumulation of virions and Env antigens on the cell surface and in turn reducing their detection by antibodies ^38,49,52^. In contrast to the conspicuous effects of Nef and Vpu deletion on V3, the exposure of V2i epitopes was not affected, consistent with our earlier findings implicating distinct mechanisms in masking V3 vs V2i epitopes ^40,45^. Nonetheless, 2219 also failed to elicit ADCC against gp120-treated CD4+ CEM.NKr cells where V3 exposure was augmented by gp120-CD4 interaction and a high binding level of 2219 to these cells was observed. In this case, the lack of ADCC activity can be attributed to 2219’s inefficient induction of Fc⍰RIIIa signaling, a requisite for effector cells to kill target cells via the ADCC mechanism. The reason for this observation remains unknown. MAbs specific for the V3 crown interact with their epitopes via the cradle or ladle binding modes, allowing different angles of approach ^24,25^. However, the binding modes did not appear to affect ADCC capability. When we tested the cradle-type V3 mAbs 2219 and 2558 and the ladle-type V3 mAb 391/95, all of which were expressed as recombinant IgG1, we detected their binding to the gp120-coated CD4+ CEM.NKr target cells but all three mAbs lacked ADCC activities.

Measurement of p24+ cells among different cells susceptible to HIV-1 infection in the humanized mouse model indicates a limitation of V3 mAb 2219 in controlling infection beyond CD4 T cells. To the best of our knowledge, our study is the first to reveal the differential potency of V3 mAb 2219 against HIV-1 infection in CD4 T cells vs monocytes. Mimicking HIV-1-infected humans, virus-infected CD34+ HSC-engrafted humanized mice harbor HIV-1 in various cell types in the blood and the lymphoid tissues ^53-55^. However, few studies evaluating neutralizing and non-neutralizing antibody functions have considered the influence of target cell types on the antibody potency. In a 2014 report Lederle et al. ^56^ demonstrated that potent bNAbs against different Env epitopes, including VRC01, PGT121, 10-1074, 2G12, b12, 4E10, and 2F5, displayed distinct inhibitory activity when pDC vs monocyte-derived DCs (moDC) were used as target cells. Interestingly, higher concentrations were consistently required for all tested mAbs to inhibit 90% infection of pDC vs moDCs target cells. Moreover, even though a relatively sensitive tier 1 HIV-1 BaL was tested, these IC90 values were much higher than those observed in the standard neutralization assay with TZM.bl target cells ^57,58^. A similar pattern was evident for non-neutralizing mAbs such as anti-gp41 mAbs 246-D and 4B3, which showed weak but detectable inhibitory activity in moDCs but not in pDCs ^56^. The mechanistic explanations for this phenomenon are yet undefined. Whether such differential resistance impacted the capacity of 2219 to control HIV-1 in CD4 T cells vs other cell types is unknown and needs further investigation. Future studies to evaluate the Env expression and Env-antibody interaction on various primary cells infected with HIV-1 are warranted.

The humanized mouse experiments in this study were designed to evaluate the prophylactic effects of V2i mAb 2158 and V3 mAb 2219 against mucosal HIV-1 exposure. Considering the absence of potent neutralizing activity, it was not surprising that these mAbs did not confer sterilizing immunity. It is worth noting, however, that the experimental system had several limitations, the most prominent being the high dose of challenge virus required to establish infection via a rectal route in this model. Two inoculations of 700 TCID50 were given to each animal based on a prior titration showing that a single inoculation yielded infection only in a fraction of the animals. This and other experimental models, e.g. the titrated multiple challenges used to infect rhesus macaques within 3 to 5 intrarectal exposures, do not reflect the transmission efficiency of HIV-1 in humans, which is estimated at a frequency of 1 to 8 transmissions per 1000 exposures ^59-61^. The challenge virus was a chimeric JRFL-NL4.3 infectious clone lacking Vpr, a viral protein important for virus spread and pathogenesis in vivo and for virus replication in myeloid cell populations ^62^. It should also be noted that we generated humanized mice using the NSG strain that lacks the C5 complement component needed to create the membrane attack complex for virion or cell lysis ^63^. Yet, independent of C5, the upstream classical complement cascade that starts from C1q binding to antibody-antigen immune complexes remains operative, producing anaphylatoxin and opsonins (C3a, C3b, iC3b, and C3d) capable of engaging G protein-coupled (C3AR1) and complement (CR1 and CR2) receptors. Indeed, the lack of virus control by the 2219 KA mutant, which is unable to bind C1q and generate C3d, implies a role of the C3 complement activity in controlling HIV-1 infection. Another caveat is that in humanized mice, there are mouse effector cells that participate in ADCP and complement-dependent functions that are not susceptible to HIV-1 infection. By contrast, in the context of non-sterilizing protection, human effector cells can be infected and, as a result, may have compromised effector functions that diminish the antibody effectiveness against the virus.

Altogether, this passive transfer study using a humanized mouse model demonstrates the ability of V2i and V3 mAbs to exert control of HIV-1 in the absence of potent neutralization and ADCC activity. Instead, ADCP and complement-mediated functions play a role in suppressing virus infection in CD4 T cells, although virus control was not achieved in other cell types such as monocytes which also harbor the virus. As V2i and V3 crown are representatives of highly immunogenic epitopes that are targeted by antibodies readily elicited by vaccination, data from this study offer further evidence that non-neutralizing activities mediated by antibodies will be important to induce with vaccines designed to prevent and control HIV-1 infection.

## Acknowledgments

This study was supported in part by NIH grant R01 AI139290 and VA Merit Review I01BX003860 to C.E.H, a CIHR foundation grant #352417 and NIH R01 AI148379 to A.F.

C.E.H. is the recipient of the US Department of Veterans Affairs BLR&D Research Career Scientist Award IK6BX004607. A.F. is the recipient of a Canada Research Chair on Retroviral Entry #RCHS0235. J.P. is the recipient of a CIHR PhD fellowship.

The authors thank Dennis Burton for the infectious molecular clone JRFL used in the ADCC assay using primary CD4+ T cells. This work was partly supported by a CIHR foundation grant #352417 and NIH R01 AI148379 to A.F.

## Materials and Methods

### Monoclonal antibodies

V2i mAb 2158 and V3 mAb 2219 were produced as recombinant IgG1 in transfected 293F cells, affinity purified by protein A (HiTrap, Sigma-Aldrich), and tested for endotoxin levels (GenScript) prior to use in experiments. For comparison, other V2i and V3 mAbs were included in some in vitro assays, whereas anti-parvovirus mAb 860-50D (designated as 860 herein) and CD4bs-specific bNAb NIH45-46 were used as negative and positive control, respectively.

The KA and LALA mutations were introduced to the Fc fragment of 2219 by QuikChange II XL Site-Directed Mutagenesis Kit (Agilent Technologies) according to the instruction manual. Like the wild type counterpart, the KA and LALA variants were produced in 293F cells following transfection.

### Humanized mouse experiments

Animal work was reviewed and approved by the University of North Carolina at Chapel Hill IACUC (ID: 17-051.0-B). Humanized mice were generated as described ^54^ by injecting human CD34+ hematopoietic stem cells (HSCs) from fetal liver tissues into the liver of irradiated NOD-SCID IL2R⍰NULL (NSG) neonates (300 rads, 0 to 2 days old, 0.5x106 cells/animal). Human fetal livers were obtained from medically indicated or elective termination of pregnancies through a non-profit intermediary working with outpatient clinics (Advanced Bioscience Resources, Alameda CA). Human leukocyte engraftment were monitored at weeks 2 and 12 by hu-CD45+ staining. Each experiment used male and female animals from the same cohort with >50% hu-CD45+ leukocytes (range of 50-66%) after week 12. The experimental protocols are outlined in Fig 1A and Fig 7A. MAb (2x 700 ug/animal) was injected intraperitoneally. Each animal was challenged rectally with 2x 700 TCID50 (equivalent to a total of 5000 pfu) of JRFL IMC, an infectious molecular clone (NFN-XS-rHSA) with chimeric JRFL-NL4.3 Vpu and Env and vpr-deleted NL4.3 backbone (a gift from Dr. J. A. Zack, UCLA) ^36^. Plasma vRNA was extracted by QIAamp Viral RNA Mini Kit (QIAGEN) and quantified by real time PCR (ABI Applied Biosystem) as in ^53^. Tissues were collected at the end of experiment, nucleic acid was extracted and cell-associated vDNA and vRNA levels were measured as described previously ^53,54,64^. Cells with active virus replication were detected by flow cytometry following mAb staining against intracellular p24 and cellular markers (CD3, CD8, CD4, CD25, FoxP3, CD14, CD11c, HLA-DR, CD123).

### ELISA with soluble Env proteins

A direct ELISA was performed as described in ^65^ to assess mAb reactivity with recombinant gp120 proteins coated on the plates. To examine mAb reactivity with virus-derived Env proteins, a sandwich ELISA was used in which 1% Trixon X-100-treated virus lysates were added to plates pre-coated with ConA (50 µg/ml, Sigma) and reacted with anti-Env mAbs ^66^. MAb binding was detected with alkaline phosphate-conjugated antibodies against human IgG or biotinylated anti-human IgG antibodies and horseradish peroxidase-streptavidin.

### Biolayer interferometry

The kinetics analysis of gp120-mAb binding was performed by biolayer interferometry using an Octet Red96 instrument (ForteBio) ^65^. mAbs were immobilized on Anti-hIgG Fc Capture (AHC) biosensors and dipped into recombinant JRFL gp120 at the designated concentrations. All samples were diluted in PBS (pH 7.4) supplemented with BSA (0.1% w/v) and Tween 20 (0.02% v/v). A baseline reference, consisting of a loaded AHC sensor run with a buffer blank for both association and dissociation steps, was utilized to correct for drift. Duplicate experiments were performed. After subtracting reference curves, data were analyzed with the Octet Data Analysis software by employing a 1:1 binding model for a global fit analysis of association and dissociation curves.

### Virus capture

MAb binding to virion was assessed by pre-incubating mAb with cell-free virus particles for 24 hours at 37oC, and treating the mAb-virus mixture with protein G-coated magnetic beads (Protein G Mag Sepharose Xtra, Cytivia). The beads were pelleted, washed, and subjected to vRNA quantification by real time PCR using the Abbott m2000 System according to the manufacturer’s instruction.

### MAb binding to Env on cells

Flow cytometry was performed to detect mAb binding to Env on transfected 293T cells ^67^ or on target cells used in the ADCC assays ^32,49,68,69^. Fluorescent secondary antibodies against human IgG were used for detection of mAb binding.

### Virus neutralization

Neutralizing activity of anti-Env mAb was measured as a reduction in β-galactosidase reporter gene expression of TZM-bl target cells. Neutralization was performed with the standard 1 hour mAb-virus incubation or the prolonged 24 hour incubation as described ^67^.

### FcγR signaling

FcγR signaling was measured according to a published protocol ^38^ using Jurkat cell–derived reporter cell lines (Jur-γRIIa and Jur-γRIIIa) that contain an integrated NFAT-driven firefly luciferase reporter gene. The Jur-γRIIa or Jur-γRIIIa cells were co-cultured for 16 hours at a 2:1 ratio with HIV-1 Δvpu-infected tetherin^high^ CD4+ lymphocytes that were pre-treated with each mAb for 15 minutes. The luciferase activity were measured using a luciferase assay kit (Promega) and subtracted with the background level obtained from co-cultures in the absence of mAb.

### ADCP

The ADCP assay was performed as described ^32^ using gp120-coated fluorescent NeutrAvidin beads and THP-1 cells or resident peritoneal macrophages from NSG mice. Beads pre-treated with mAbs were added to THP-1 cells, and phagocytosis was measured by flow cytometry after an overnight incubation. ADCP scores were calculated as: (percentage of bead-positive cells⍰×⍰mean fluorescence intensity of bead-positive cells)/10^5^.

### Complement deposition

C1q binding to immune complexes made with V2i mAb 2159 or V3 mAb 2219 was measured in ELISA. MAb was reacted with antigen coated on the plates and then treated with serially diluted C1q from human serum (Sigma). C1q binding was detected with horseradish peroxidase-conjugated anti-C1q antibody. C3d deposition was detected using Luminex assay according to a published protocol ^70,71^ with some modifications. Antigen-coupled xMAP beads were incubated with serially diluted mAb, and then treated with human complement serum (33.3%, Sigma) at 37oC for 1 hour. C3d production and deposition was measured by biotinylated anti-C3d mAb (Quidel) and PE-streptavidin.

### ADCC

The ADCC assay using gp120-coated CD4+ CEM.NKr target cells and PBMCs as effector cells was done as described in ^68^, whereas the assay with virus-infected NKR24 reporter cells and human NK cell line KHYG-1 was done according to ^69^. The third ADCC assay was performed using full length HIV-1 JRFL IMC-infected primary CD4 T cells and PBMC effector cells ^49,72^.

## References

1 UNAIDS. Global HIV & AIDS statistics - 2020 fact sheet, <https://www.unaids.org/en/resources/fact-sheet> (2020).

2 Rerks-Ngarm, S. et al. Vaccination with ALVAC and AIDSVAX to prevent HIV-1 infection in Thailand. N Engl J Med 361, 2209–2220, doi:10.1056/NEJMoa0908492 (2009).

3 Haynes, B. F. et al. Immune-correlates analysis of an HIV-1 vaccine efficacy trial. N Engl J Med 366, 1275–1286, doi:10.1056/NEJMoa1113425 (2012).

4 Kim, J. H., Excler, J. L. & Michael, N. L. Lessons from the RV144 Thai phase III HIV-1 vaccine trial and the search for correlates of protection. Annu Rev Med 66, 423–437, doi:10.1146/annurev-med-052912-123749 (2015).

5 Zolla-Pazner, S. et al. Analysis of V2 antibody responses induced in vaccinees in the ALVAC/AIDSVAX HIV-1 vaccine efficacy trial. PLoS One 8, e53629, doi:10.1371/journal.pone.0053629 (2013).

6 Gottardo, R. et al. Plasma IgG to linear epitopes in the V2 and V3 regions of HIV-1 gp120 correlate with a reduced risk of infection in the RV144 vaccine efficacy trial. PLoS One 8, e75665, doi:10.1371/journal.pone.0075665 (2013).

7 Zolla-Pazner, S. et al. Vaccine-induced Human Antibodies Specific for the Third Variable Region of HIV-1 gp120 Impose Immune Pressure on Infecting Viruses. EBioMedicine 1, 37–45, doi:10.1016/j.ebiom.2014.10.022 (2014).

8 Bonsignori, M. et al. Antibody-dependent cellular cytotoxicity-mediating antibodies from an HIV-1 vaccine efficacy trial target multiple epitopes and preferentially use the VH1 gene family. J Virol 86, 11521–11532, doi:10.1128/JVI.01023-12 (2012).

9 Burton, D. R. et al. Limited or no protection by weakly or nonneutralizing antibodies against vaginal SHIV challenge of macaques compared with a strongly neutralizing antibody. Proc Natl Acad Sci U S A 108, 11181–11186, doi:10.1073/pnas.1103012108 (2011).

10 Oliphant, T. et al. Antibody recognition and neutralization determinants on domains I and II of West Nile Virus envelope protein. J Virol 80, 12149–12159, doi:10.1128/JVI.01732-06 (2006).

11 Vogt, M. R. et al. Poorly neutralizing cross-reactive antibodies against the fusion loop of West Nile virus envelope protein protect in vivo via Fcgamma receptor and complement-dependent effector mechanisms. J Virol 85, 11567–11580, doi:10.1128/JVI.05859-11 (2011).

12 Kim, J. H. et al. Non-neutralizing antibodies induced by seasonal influenza vaccine prevent, not exacerbate A(H1N1)pdm09 disease. Sci Rep 6, 37341, doi:10.1038/srep37341 (2016).

13 Tan, G. S. et al. Broadly-Reactive Neutralizing and Non-neutralizing Antibodies Directed against the H7 Influenza Virus Hemagglutinin Reveal Divergent Mechanisms of Protection. PLoS Pathog 12, e1005578, doi:10.1371/journal.ppat.1005578 (2016).

14 DiLillo, D. J., Palese, P., Wilson, P. C. & Ravetch, J. V. Broadly neutralizing anti-influenza antibodies require Fc receptor engagement for in vivo protection. J Clin Invest 126, 605–610, doi:10.1172/JCI84428 (2016).

15 Carragher, D. M., Kaminski, D. A., Moquin, A., Hartson, L. & Randall, T. D. A novel role for non-neutralizing antibodies against nucleoprotein in facilitating resistance to influenza virus. J Immunol 181, 4168–4176, doi:10.4049/jimmunol.181.6.4168 (2008).

16 Santra, S. et al. Human Non-neutralizing HIV-1 Envelope Monoclonal Antibodies Limit the Number of Founder Viruses during SHIV Mucosal Infection in Rhesus Macaques. PLoS Pathog 11, e1005042, doi:10.1371/journal.ppat.1005042 (2015).

17 Horwitz, J. A. et al. Non-neutralizing Antibodies Alter the Course of HIV-1 Infection In Vivo. Cell 170, 637–648 e610, doi:10.1016/j.cell.2017.06.048 (2017).

18 Eda, Y. et al. Anti-V3 humanized antibody KD-247 effectively suppresses ex vivo generation of human immunodeficiency virus type 1 and affords sterile protection of monkeys against a heterologous simian/human immunodeficiency virus infection. J Virol 80, 5563–5570, doi:10.1128/JVI.02095-05 (2006).

19 Hessell, A. J. et al. Reduced Cell-Associated DNA and Improved Viral Control in Macaques following Passive Transfer of a Single Anti-V2 Monoclonal Antibody and Repeated Simian/Human Immunodeficiency Virus Challenges. J Virol 92, doi:10.1128/JVI.02198-17 (2018).

20 Pan, R., Gorny, M. K., Zolla-Pazner, S. & Kong, X. P. The V1V2 Region of HIV-1 gp120 Forms a Five-Stranded Beta Barrel. J Virol 89, 8003–8010, doi:10.1128/JVI.00754-15 (2015).

21 Mayr, L. M., Cohen, S., Spurrier, B., Kong, X. P. & Zolla-Pazner, S. Epitope mapping of conformational V2-specific anti-HIV human monoclonal antibodies reveals an immunodominant site in V2. PLoS One 8, e70859, doi:10.1371/journal.pone.0070859 (2013).

22 Gorny, M. K. et al. Functional and immunochemical cross-reactivity of V2-specific monoclonal antibodies from HIV-1-infected individuals. Virology 427, 198–207, doi:10.1016/j.virol.2012.02.003 (2012).

23 Spurrier, B., Sampson, J., Gorny, M. K., Zolla-Pazner, S. & Kong, X. P. Functional implications of the binding mode of a human conformation-dependent V2 monoclonal antibody against HIV. J Virol 88, 4100–4112, doi:10.1128/JVI.03153-13 (2014).

24 Balasubramanian, P. et al. Differential induction of anti-V3 crown antibodies with cradle-and ladle-binding modes in response to HIV-1 envelope vaccination. Vaccine 35, 1464–1473, doi:10.1016/j.vaccine.2016.11.107 (2017).

25 Jiang, X. et al. Conserved structural elements in the V3 crown of HIV-1 gp120. Nat Struct Mol Biol 17, 955–961, doi:10.1038/nsmb.1861 (2010).

26 Stanfield, R. L., Gorny, M. K., Zolla-Pazner, S. & Wilson, I. A. Crystal structures of human immunodeficiency virus type 1 (HIV-1) neutralizing antibody 2219 in complex with three different V3 peptides reveal a new binding mode for HIV-1 cross-reactivity. J Virol 80, 6093–6105, doi:10.1128/JVI.00205-06 (2006).

27 Li, L. et al. A broad range of mutations in HIV-1 neutralizing human monoclonal antibodies specific for V2, V3, and the CD4 binding site. Mol Immunol 66, 364–374, doi:10.1016/j.molimm.2015.04.011 (2015).

28 Hioe, C. E. et al. Anti-V3 monoclonal antibodies display broad neutralizing activities against multiple HIV-1 subtypes. PLoS One 5, e10254, doi:10.1371/journal.pone.0010254 (2010).

29 Gorny, M. K. et al. Cross-clade neutralizing activity of human anti-V3 monoclonal antibodies derived from the cells of individuals infected with non-B clades of human immunodeficiency virus type 1. J Virol 80, 6865–6872, doi:10.1128/JVI.02202-05 (2006).

30 Gorny, M. K. et al. Human monoclonal antibodies specific for conformation-sensitive epitopes of V3 neutralize human immunodeficiency virus type 1 primary isolates from various clades. J Virol 76, 9035–9045, doi:10.1128/jvi.76.18.9035-9045.2002 (2002).

31 Agarwal, A., Hioe, C. E., Swetnam, J., Zolla-Pazner, S. & Cardozo, T. Quantitative assessment of masking of neutralization epitopes in HIV-1. Vaccine 29, 6736–6741, doi:10.1016/j.vaccine.2010.12.052 (2011).

32 Balasubramanian, P. et al. Functional Antibody Response Against V1V2 and V3 of HIV gp120 in the VAX003 and VAX004 Vaccine Trials. Sci Rep 8, 542, doi:10.1038/s41598-017-18863-0 (2018).

33 Zolla-Pazner, S. et al. Vaccine-induced IgG antibodies to V1V2 regions of multiple HIV-1 subtypes correlate with decreased risk of HIV-1 infection. PLoS One 9, e87572, doi:10.1371/journal.pone.0087572 (2014).

34 Karasavvas, N. et al. The Thai Phase III HIV Type 1 Vaccine trial (RV144) regimen induces antibodies that target conserved regions within the V2 loop of gp120. AIDS Res Hum Retroviruses 28, 1444–1457, doi:10.1089/aid.2012.0103 (2012).

35 Corey, L. et al. Immune correlates of vaccine protection against HIV-1 acquisition. Sci Transl Med 7, 310rv317, doi:10.1126/scitranslmed.aac7732 (2015).

36 Roth, M. D. et al. Cocaine enhances human immunodeficiency virus replication in a model of severe combined immunodeficient mice implanted with human peripheral blood leukocytes. J Infect Dis 185, 701–705, doi:10.1086/339012 (2002).

37 Gigler, A. et al. Generation of neutralizing human monoclonal antibodies against parvovirus B19 proteins. J Virol 73, 1974–1979, doi:10.1128/JVI.73.3.1974-1979.1999 (1999).

38 Alvarez, R. A. et al. Enhanced FCGR2A and FCGR3A signaling by HIV viremic controller IgG. JCI Insight 2, e88226, doi:10.1172/jci.insight.88226 (2017).

39 Binley, J. M. et al. Comprehensive cross-clade neutralization analysis of a panel of anti-human immunodeficiency virus type 1 monoclonal antibodies. J Virol 78, 13232–13252, doi:10.1128/JVI.78.23.13232-13252.2004 (2004).

40 Upadhyay, C. et al. Distinct mechanisms regulate exposure of neutralizing epitopes in the V2 and V3 loops of HIV-1 envelope. J Virol 88, 12853–12865, doi:10.1128/JVI.02125-14 (2014).

41 Tay, M. Z., Wiehe, K. & Pollara, J. Antibody-Dependent Cellular Phagocytosis in Antiviral Immune Responses. Front Immunol 10, 332, doi:10.3389/fimmu.2019.00332 (2019).

42 Ackerman, M. E. et al. A robust, high-throughput assay to determine the phagocytic activity of clinical antibody samples. J Immunol Methods 366, 8–19, doi:10.1016/j.jim.2010.12.016 (2011).

43 Forthal, D. N. & Finzi, A. Antibody-dependent cellular cytotoxicity in HIV infection. AIDS 32, 2439–2451, doi:10.1097/QAD.0000000000002011 (2018).

44 Richard, J., Prevost, J., Alsahafi, N., Ding, S. & Finzi, A. Impact of HIV-1 Envelope Conformation on ADCC Responses. Trends Microbiol 26, 253–265, doi:10.1016/j.tim.2017.10.007 (2018).

45 Powell, R. L. R. et al. Plasticity and Epitope Exposure of the HIV-1 Envelope Trimer. J Virol 91, doi:10.1128/JVI.00410-17 (2017).

46 Hezareh, M., Hessell, A. J., Jensen, R. C., van de Winkel, J. G. & Parren, P. W. Effector function activities of a panel of mutants of a broadly neutralizing antibody against human immunodeficiency virus type 1. J Virol 75, 12161–12168, doi:10.1128/JVI.75.24.12161-12168.2001 (2001).

47 Chung, A. W. et al. Identification of antibody glycosylation structures that predict monoclonal antibody Fc-effector function. AIDS 28, 2523–2530, doi:10.1097/QAD.0000000000000444 (2014).

48 Veillette, M. et al. Interaction with cellular CD4 exposes HIV-1 envelope epitopes targeted by antibody-dependent cell-mediated cytotoxicity. J Virol 88, 2633–2644, doi:10.1128/JVI.03230-13 (2014).

49 Veillette, M. et al. The HIV-1 gp120 CD4-bound conformation is preferentially targeted by antibody-dependent cellular cytotoxicity-mediating antibodies in sera from HIV-1-infected individuals. J Virol 89, 545–551, doi:10.1128/JVI.02868-14 (2015).

50 Alsahafi, N. et al. Nef Proteins from HIV-1 Elite Controllers Are Inefficient at Preventing Antibody-Dependent Cellular Cytotoxicity. J Virol 90, 2993–3002, doi:10.1128/JVI.02973-15 (2015).

51 Prevost, J. et al. Incomplete Downregulation of CD4 Expression Affects HIV-1 Env Conformation and Antibody-Dependent Cellular Cytotoxicity Responses. J Virol 92, doi:10.1128/JVI.00484-18 (2018).

52 Arias, J. F. et al. Tetherin antagonism by Vpu protects HIV-infected cells from antibody-dependent cell-mediated cytotoxicity. Proc Natl Acad Sci U S A 111, 6425–6430, doi:10.1073/pnas.1321507111 (2014).

53 Li, G. et al. Regulatory T Cells Contribute to HIV-1 Reservoir Persistence in CD4+ T Cells Through Cyclic Adenosine Monophosphate-Dependent Mechanisms in Humanized Mice In Vivo. J Infect Dis 216, 1579–1591, doi:10.1093/infdis/jix547 (2017).

54 Li, G. et al. HIV-1 infection depletes human CD34+CD38-hematopoietic progenitor cells via pDC-dependent mechanisms. PLoS Pathog 13, e1006505, doi:10.1371/journal.ppat.1006505 (2017).

55 Zhao, J. et al. Infection and depletion of CD4+ group-1 innate lymphoid cells by HIV-1 via type-I interferon pathway. PLoS Pathog 14, e1006819, doi:10.1371/journal.ppat.1006819 (2018).

56 Lederle, A. et al. Neutralizing antibodies inhibit HIV-1 infection of plasmacytoid dendritic cells by an FcgammaRIIa independent mechanism and do not diminish cytokines production. Sci Rep 4, 5845, doi:10.1038/srep05845 (2014).

57 Lorenzi, J. C. C. et al. Neutralizing Activity of Broadly Neutralizing anti-HIV-1 Antibodies against Primary African Isolates. J Virol, doi:10.1128/JVI.01909-20 (2020).

58 Cohen, Y. Z. et al. Neutralizing Activity of Broadly Neutralizing Anti-HIV-1 Antibodies against Clade B Clinical Isolates Produced in Peripheral Blood Mononuclear Cells. J Virol 92, doi:10.1128/JVI.01883-17 (2018).

59 Wawer, M. J. et al. Rates of HIV-1 transmission per coital act, by stage of HIV-1 infection, in Rakai, Uganda. J Infect Dis 191, 1403–1409, doi:10.1086/429411 (2005).

60 Hollingsworth, T. D., Anderson, R. M. & Fraser, C. HIV-1 transmission, by stage of infection. J Infect Dis 198, 687–693, doi:10.1086/590501 (2008).

61 Hughes, J. P. et al. Determinants of per-coital-act HIV-1 infectivity among African HIV-1-serodiscordant couples. J Infect Dis 205, 358–365, doi:10.1093/infdis/jir747 (2012).

62 Nodder, S. B. & Gummuluru, S. Illuminating the Role of Vpr in HIV Infection of Myeloid Cells. Front Immunol 10, 1606, doi:10.3389/fimmu.2019.01606 (2019).

63 Baxter, A. G. & Cooke, A. Complement lytic activity has no role in the pathogenesis of autoimmune diabetes in NOD mice. Diabetes 42, 1574–1578, doi:10.2337/diab.42.11.1574 (1993).

64 Cheng, L., Ma, J., Li, G. & Su, L. Humanized Mice Engrafted With Human HSC Only or HSC and Thymus Support Comparable HIV-1 Replication, Immunopathology, and Responses to ART and Immune Therapy. Front Immunol 9, 817, doi:10.3389/fimmu.2018.00817 (2018).

65 Hioe, C. E. et al. Modulation of Antibody Responses to the V1V2 and V3 Regions of HIV-1 Envelope by Immune Complex Vaccines. Front Immunol 9, 2441, doi:10.3389/fimmu.2018.02441 (2018).

66 Wang, S. et al. Polyvalent HIV-1 Env vaccine formulations delivered by the DNA priming plus protein boosting approach are effective in generating neutralizing antibodies against primary human immunodeficiency virus type 1 isolates from subtypes A, B, C, D and E. Virology 350, 34–47, doi:10.1016/j.virol.2006.02.032 (2006).

67 Upadhyay, C. et al. Signal peptide of HIV-1 envelope modulates glycosylation impacting exposure of V1V2 and other epitopes. PLoS Pathog 16, e1009185, doi:10.1371/journal.ppat.1009185 (2020).

68 Costa, M. R. et al. Fc Receptor-Mediated Activities of Env-Specific Human Monoclonal Antibodies Generated from Volunteers Receiving the DNA Prime-Protein Boost HIV Vaccine DP6-001. J Virol 90, 10362–10378, doi:10.1128/JVI.01458-16 (2016).

69 Hessell, A. J. et al. Virus Control in Vaccinated Rhesus Macaques Is Associated with Neutralizing and Capturing Antibodies against the SHIV Challenge Virus but Not with V1V2 Vaccine-Induced Anti-V2 Antibodies Alone. J Immunol 206, 1266–1283, doi:10.4049/jimmunol.2001010 (2021).

70 Weiss, S. et al. A High-Throughput Assay for Circulating Antibodies Directed Against the S Protein of Severe Acute Respiratory Syndrome Coronavirus 2. J Infect Dis 222, 1629–1634, doi:10.1093/infdis/jiaa531 (2020).

71 Perez, L. G. et al. V1V2-specific complement activating serum IgG as a correlate of reduced HIV-1 infection risk in RV144. PLoS One 12, e0180720, doi:10.1371/journal.pone.0180720 (2017).

72 Alsahafi, N. et al. An Asymmetric Opening of HIV-1 Envelope Mediates Antibody-Dependent Cellular Cytotoxicity. Cell Host Microbe 25, 578–587 e575, doi:10.1016/j.chom.2019.03.002 (2019).

