## Supplemental Figures for "Non-neutralizing antibodies targeting the immunogenic regions of HIV-1 envelope reduce mucosal infection and virus burden in humanized mice"

Figure S1

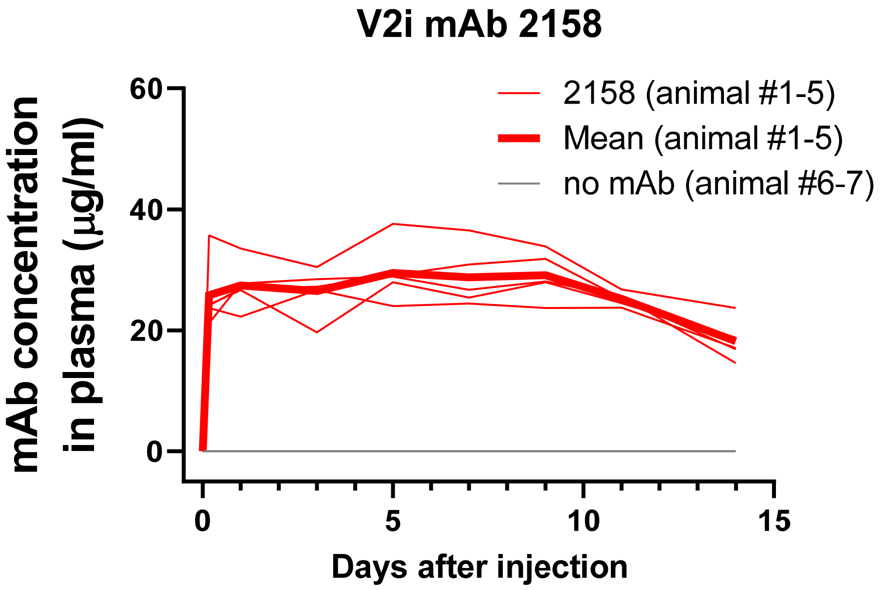


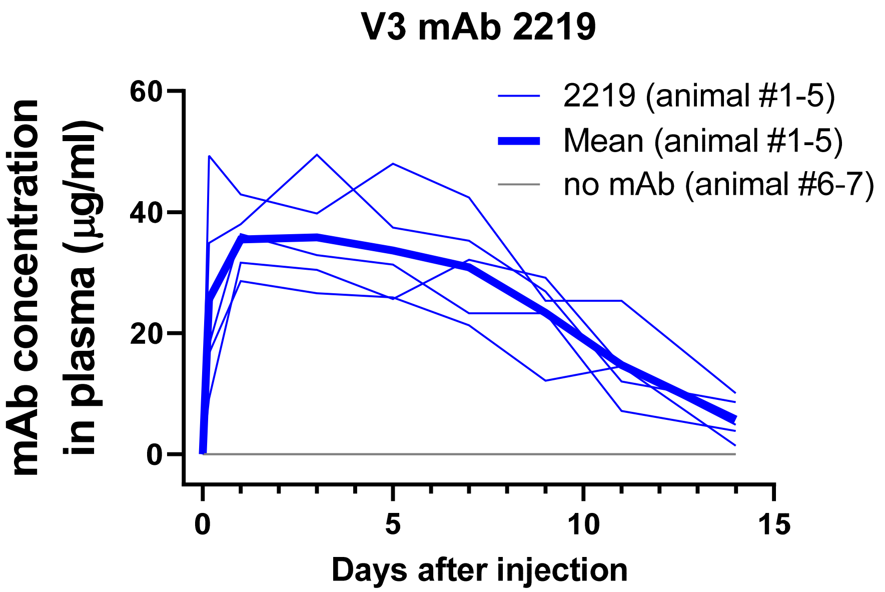


**Supplemental Figure S1: Pharmacokinetic study of human V2i mAb 2158 and V3 mAb 2219 in NSG mice.**

Each mice was given V2i mAb 2158 and V3 mAb 2219 (700 μg per mAb) intraperitoneally. Plasma was collected and monitored for the concentration of V2i mAb 2158 and V3 mAb 2219 from day 0 to day 14 using ELISA with V1V2-1FD6 or peptide V3 antigens and the respective mAbs as standard. Based on the average values, the half life was estimated to be >14 days for V2i mAb 2158 and 11 days for V3 mAb 2219.

Figure S2

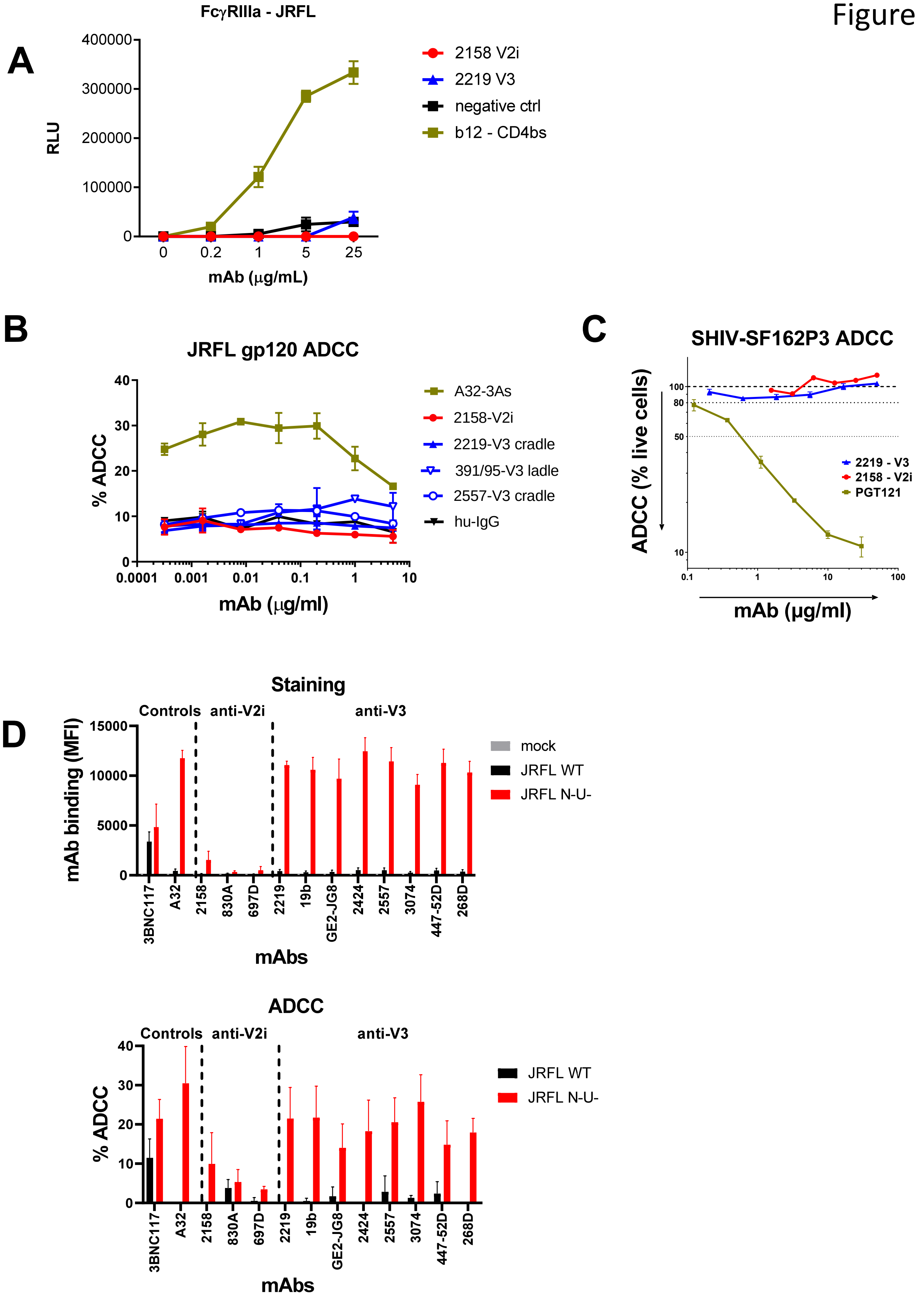
A
FcγRIIIa - JRFL
RLU
400000
300000
200000
100000
0
0
0.2
1
5
25
mAb (μg/mL)
2158 V2i
2219 V3
negative ctrl
b12 - CD4bs
B
JRFL gp120 ADCC
% ADCC
40
30
20
10
0
0.0001
0.001
0.01
0.1
1
10
mAb (μg/ml)
A32-3As
2158-V2i
2219-V3 cradle
391/95-V3 ladle
2557-V3 cradle
hu-IgG
C
SHIV-SF162P3 ADCC
ADCC (% live cells)
100
80
50
10
0.1
1
10
100
mAb (μg/ml)
2219 - V3
2158 - V2i
PGT121
D
Staining
Controls
anti-V2i
anti-V3
mAb binding (MFI)
15000
10000
5000
0
3BNC117
A32
2158
830A
697D
2219
19b
GE2-JG8
2424
2557
3074
447-52D
268D
mAbs
mock
JRFL WT
JRFL N-U-
ADCC
Controls
anti-V2i
anti-V3
% ADCC
40
30
20
10
0
3BNC117
A32
2158
830A
697D
2219
19b
GE2-JG8
2424
2557
3074
447-52D
268D
mAbs
JRFL WT
JRFL N-U-

**Supplemental Figure S2:**
**V2i mAb 2158 and V3 mAb 2219 lack the ability to mediate FcγRIIIa signaling and ADCC activity.**

A) FcγRIIIA signaling was measured by co-incubating JRFL Δvpu-nucleofected Jurkat cells with Jur-γRIIIa luciferase reporter cells in the presence of V2i, V3, or control mAbs. CD4-binding site mAb b12 served as a positive control. RLU: relative light unit.

B) ADCC activity was determined using $CD4^+$ CEM.NKr target cells that were coated with recombinant gp120 JRFL, treated with V2i, V3, or control mAbs, and incubated with PBMCs as effector cells. V2i mAb, cradle-type V3 mAbs 2219 and 2557, and ladle-type V3 mAb 391/95 were tested along with anti-C1C2 mAb A32-3As (positive control) and purified human IgG (negative control).

C) ADCC activity was examined in a second assay in which NKR24 luciferase reporter cells were infected with virus for 3 to 4 days, combined with the effector cells, human $CD16^+$ NK cell line KHYG-1, at an effector-to-target cell ratio of 5:1, and incubated with serially diluted mAbs for 8 hours. The NKR24 target cell viability was measured by luciferase activity. PGT121 was used as a positive control.

D) ADCC activity was also determined against primary CD4+ T cells infected with JRFL IMC or the Nef and Vpu-deleted counterpart. Binding of 2158 and 2219 to virus- vs mock-infected target cells was first examined by flow cytometry (top). Other V2i and V3 mAbs were tested for comparison. 3BNC117 (anti-CD4bs) and A32 (anti-C1C2) served as positive controls. ADCC were subsequently measured with PBMC effector cells at an effector-to-target ratio of 10:1 (bottom). Viability dye was used to measure cytotoxicity against infected target cells. Each mAb was tested at 5 μg/mL.

Figure S3

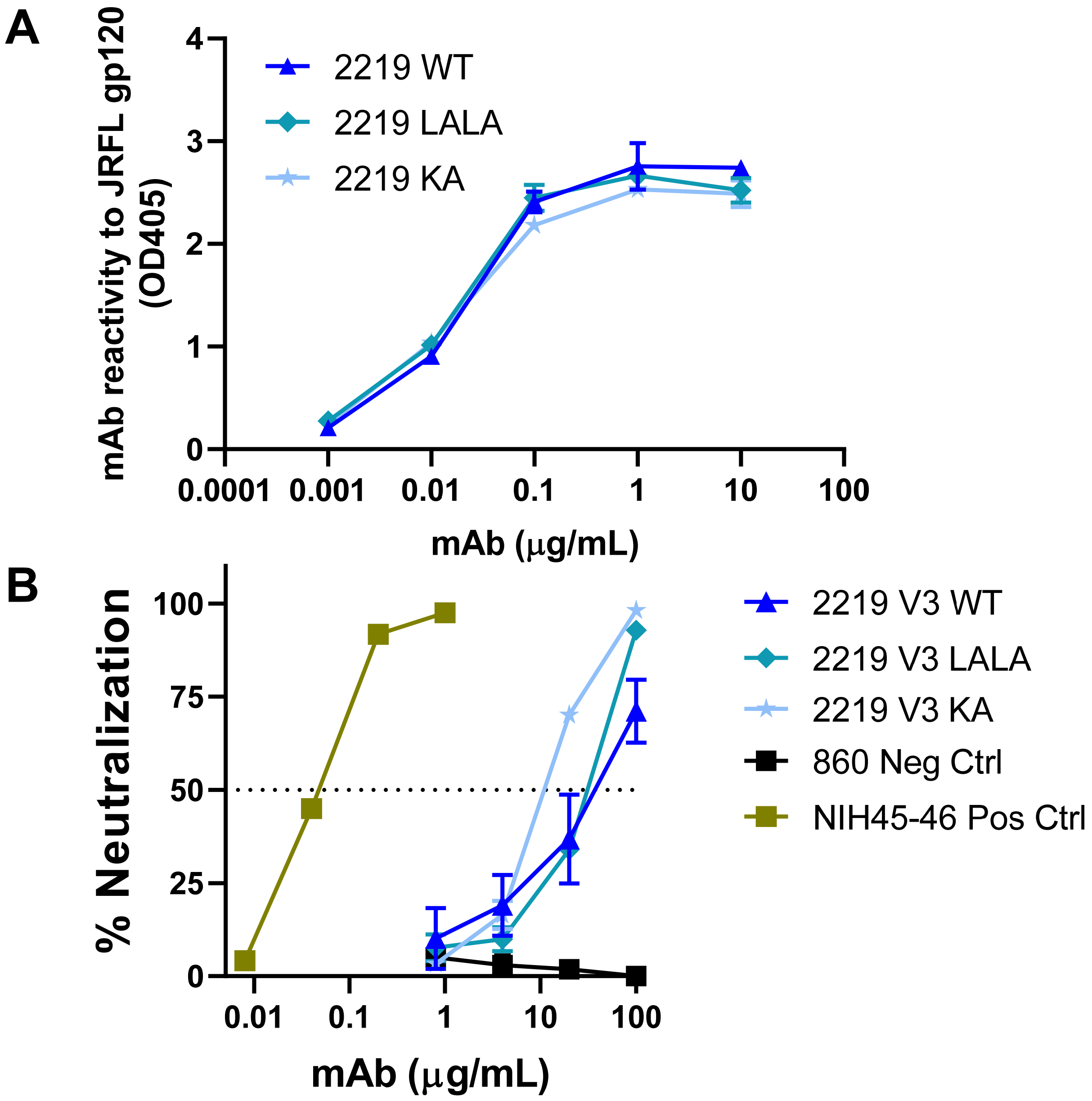


**Figure S3: Fc mutations LALA and KA do not alter gp120 binding or neutralizing activity of V3 mAb 2219.**

A) ELISA reactivity of 2219 WT vs Fc mutants was tested against recombinant gp120 JRFL.

B) Neutralization of JRFL by 2219 WT vs Fc mutants was assessed with TZM.bl target cells after 24-hour mAb-virus incubation. CD4bs-specific bNAb NIH45-46 and irrelevant mAb 860 were included as controls.

Figure S4

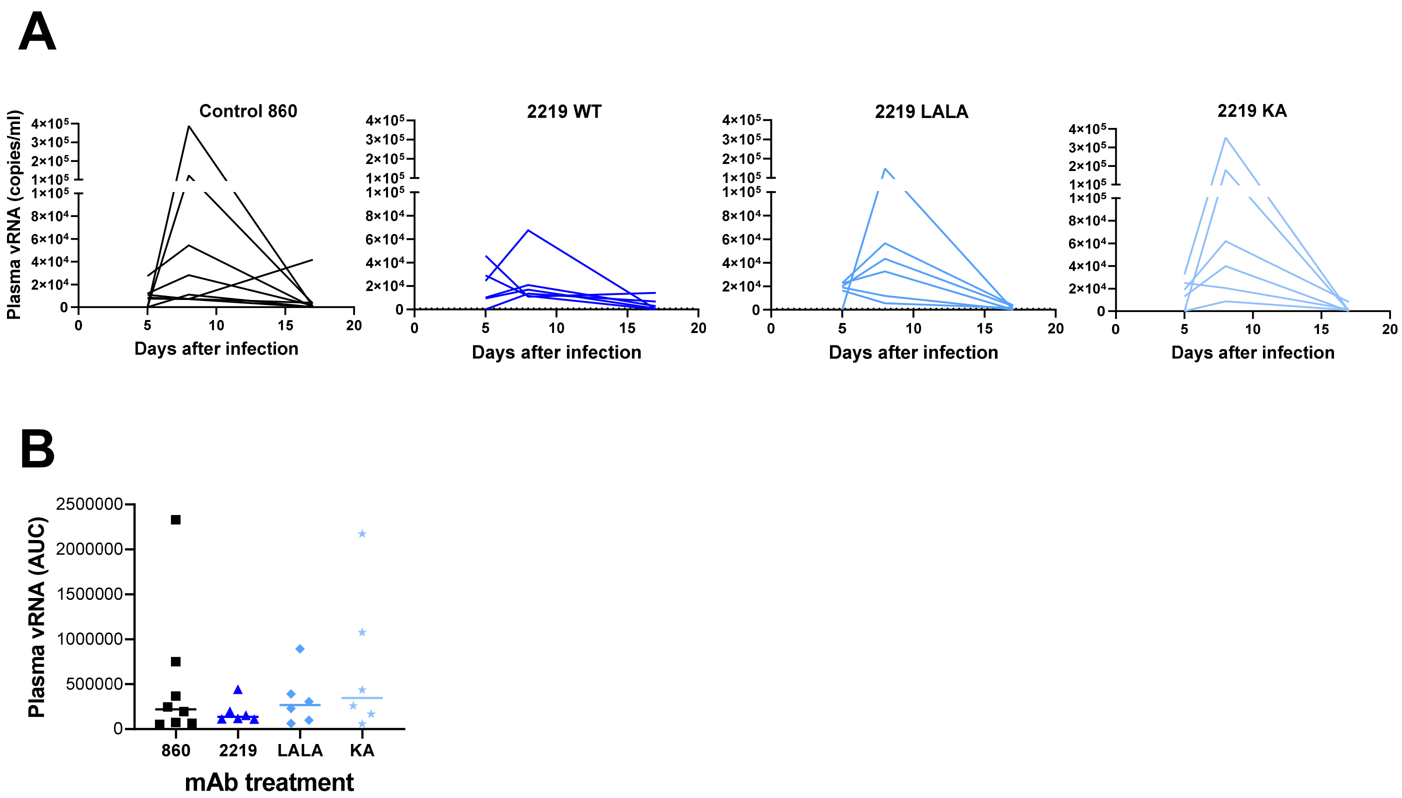


**Figure S4. Plasma vRNA load in humanized mice treated with anti-V3 mAb 2219 WT or Fc mutants and challenged with JRFL IMC.**
Another set of experiment was performed in which humanized animals were given WT, LALA, or KA 2219 mAbs intraperitoneally and then challenged with HIV-1 JRFL rectally following the protocol in Fig 1A, except that blood was collected on day 5, day 8, and day 17.

A. Virus load in plasma was measured over time after infection.
B. AUC of plasma vRNA in individual mice from day 5 to day 17.
